# Self-assembly of progenitor cells under the aegis of platelet factors facilitates human skin organoid formation and vascularized wound healing

**DOI:** 10.1101/2020.09.10.292409

**Authors:** Patricia Peking, Linda Krisch, Martin Wolf, Anna Hoog, Balázs Vári, Katharina Muigg, Rodolphe Poupardin, Cornelia Scharler, Elisabeth Russe, Harald Stachelscheid, Achim Schneeberger, Katharina Schallmoser, Dirk Strunk

## Abstract

Stem/progenitor cells can self-organize into organoids modelling tissue function and regeneration. Here we demonstrate that human platelet-derived factors can orchestrate 3D self-assembly of clonally expanded adult skin fibroblasts, keratinocytes and endothelial progenitors forming skin organoids within three days. Organoids showed distinct signaling patterns in response to inflammatory stimuli that clearly differed from separated cell types. Human induced pluripotent stem cell (hiPSC)-derived skin cell progenitors also self-assembled into stratified human skin within two weeks, healing deep wounds of immune-deficient mice. Co-transplantation of endothelial progenitors significantly accelerated vascularization. Mechanistically, platelet-derived extracellular vesicles mediated the platelet-derived trophic effects. Long-term fitness of epidermal cells was accelerated further by keratinocyte growth factor mRNA transfection. No tumorigenesis was observed upon xenografting. This permits novel rapid 3D skin-related pharmaceutical testing opportunities and facilitates development of iPSC-based skin regeneration strategies.

---

The human skin is the body’s largest organ, fulfilling multiple biochemical and sensory functions in addition to forming the physical body boundary. Locally damaged adult skin can heal with scars but lacks the capacity to fully regenerate the complex composition of larger skin areas ^1^. Severe scarring and chronic wounds after major skin burns ^2^, surgeries or devastating skin diseases ^3^ limit skin mobility, respiration and light protection, eventually resulting in body fluid loss, life-threatening infections and skin cancer ^4,5^.

The current gold standard for extended area epidermal replacement is transplantation of ex vivo engineered epidermal sheets ^6,7^. Improved dermal restoration and vascularization strategies are prerequisite for providing an oxygen-rich environment that promotes extended skin tissue regeneration ^8^. In vitro generated dermis equivalents with pre-formed vessels showed enhanced full-thickness skin wound repair in rats ^9^. Transplantation of endothelial cells could induce neo-vascularization upon co-transplantation of stromal cells that acted as pericytes stabilizing de-novo formed vessels and reducing endothelial immunogenicity ^10–12^. Platelet-derived regenerative factors promoted angiogenesis, collagen synthesis and epithelialization ^13,14^. The use of hiPSCs for tissue regeneration has a great potential due to their proliferative potential generating virtually unlimited cell amounts including somatic as well as stromal and endothelial cells ^15^. Various cell types can be generated from a single hiPSC clone, allowing the reconstruction of complex organs.

For many other organs, stem/progenitor cell-derived 3D organoids have been established as a tool for biomedical research and development of cell-based organ transplantation or regeneration strategies. Appropriate vascularization was considered to better mimic complex organ architecture in organoid test systems and perhaps even create instantly transplantable tissue ^16–18^. Conventional layered skin equivalents, though 3D, are not necessarily considered to represent a predecessor of current spheroid organoids. First described in 1980s based on pioneering co-cultures of keratinocytes on fibroblasts at the air-liquid interface, basically established by Rheinwald and Green ^19^, refined skin equivalents are meanwhile validated and licensed tools, upon 5-6-week manufacturing, for basic research and toxicity testing, replacing animal experimentation as part of global 3R efforts ^20^. Most recently, complex organoids containing hairy skin with appendages recapitulating second trimester human tissue were generated from hiPSCs with a sophisticated protocol over 4-5 months incubation periods. After removing subcutaneous cartilage and other extracutaneous structures, more than half of the organoids even established hairy human skin on mice opening entirely new opportunities to study human skin development ^21^.

Extended time to create transplantable skin equivalents, missing pre-vascularization and lack of superior outcome in clinical testing compared to invasively obtained split skin transplants have hampered broader clinical applicability ^22^. Self-organization of cells into three-dimensional multicellular organoid-like structures may be enhanced by advanced bioengineering strategies including 3D printing ^23^. Towards scar-free skin regeneration, various improvements still need to be achieved to get regionalized skin comprizing loco-typic hairiness and appendages ideally also enabling creation of perfect organoid model systems to read-out complex transcriptional regulatory networks in their response to environmental or pharmacologic input signals ^24,25^. It remains to be clarified to what extent the rather simplistic organoids can facilitate intercellular communication analysis and development of novel pharmacologic or diagnostic strategies ^26^.

Here we introduce a dual strategy essentially based on rapid skin progenitor self-assembly into (1) spheroid vascularized organoids in vitro and (2) properly stratified rapidly vascularized human skin in vivo. The complex mixture of platelet-derived growth and regeneration factors in human platelet lysate (hPL) acted in a dose-dependent manner to orchestrate self-organization and de novo transplant vasculogenesis, virtually mimicking wound repair processes^13^ in vitro as well as, after co-transplantation of human endothelial cells with fibroblasts and keratinocytes, in vivo. Comparative molecular response profiling showed distinct signaling patterns in organoids that clearly differed from separated cell types. Both adult progenitor cell types and iPSC-derived matured skin cells were effective for healing deep skin wounds, thus extending applicability of cell transplants.

## Results

### Adult progenitor cell isolation, characterization and large scale expansion

Primary keratinocytes and fibroblasts were isolated from epidermal and dermal split-skin parts, respectively, and endothelial colony-forming progenitor cells (ECFCs) as a source of high endothelial cell numbers based on previously established protocols by late outgrowth from umbilical cord blood ^11,27,28^. All cell types were large-scale expanded under animal serum-free conditions ^29,30^ (**Fig. S1a**). Cell characterization showed specific CD90^+^ fibroblasts, keratin 14^+^ keratinocytes and CD31^+^ ECFCs (**Fig. 1ab**). Clonogenicity indicated progenitor enrichment after low seeding density propagation selecting for high proliferative potential cells ^11,31,32^ and showing donor-dependent colony-forming unit (CFU) capacities ranging from 42-140% in fibroblasts and 60-100% in ECFC preparations, respectively (**Fig. 1c**). Keratinocytes grew clonally on feeders after primary culture (**Fig. 1d**) with donor variation in morphology and keratin 14 expression between 19.1-97.2% (**Fig. S1b**). More than 90% pure keratin 14^+^ cultures, enriched for highly proliferative basal epidermal keratinocyte progenitors, were selected for subsequent organoid formation and transplantation experiments. ECFC functionality was confirmed by vascular network formation (**Fig. 1e**). Because progenitor cells from many other organs can assemble in spheroids when subjected to ultralow attachment environment, we tested this behavior for the three cell types. Both fibroblasts and keratinocytes regularly formed compact 3D spheroids. Cord blood-derived ECFCs formed 2D colonies (**Fig. 1c**) and loosely accumulated in 3D (**Fig. 1f**). These results prompted us to ask the question if it is possible to instantly create spheroid skin organoids using contemporary technology clearly different from protracted conventional skin equivalent protocols.

**Figure 1:**
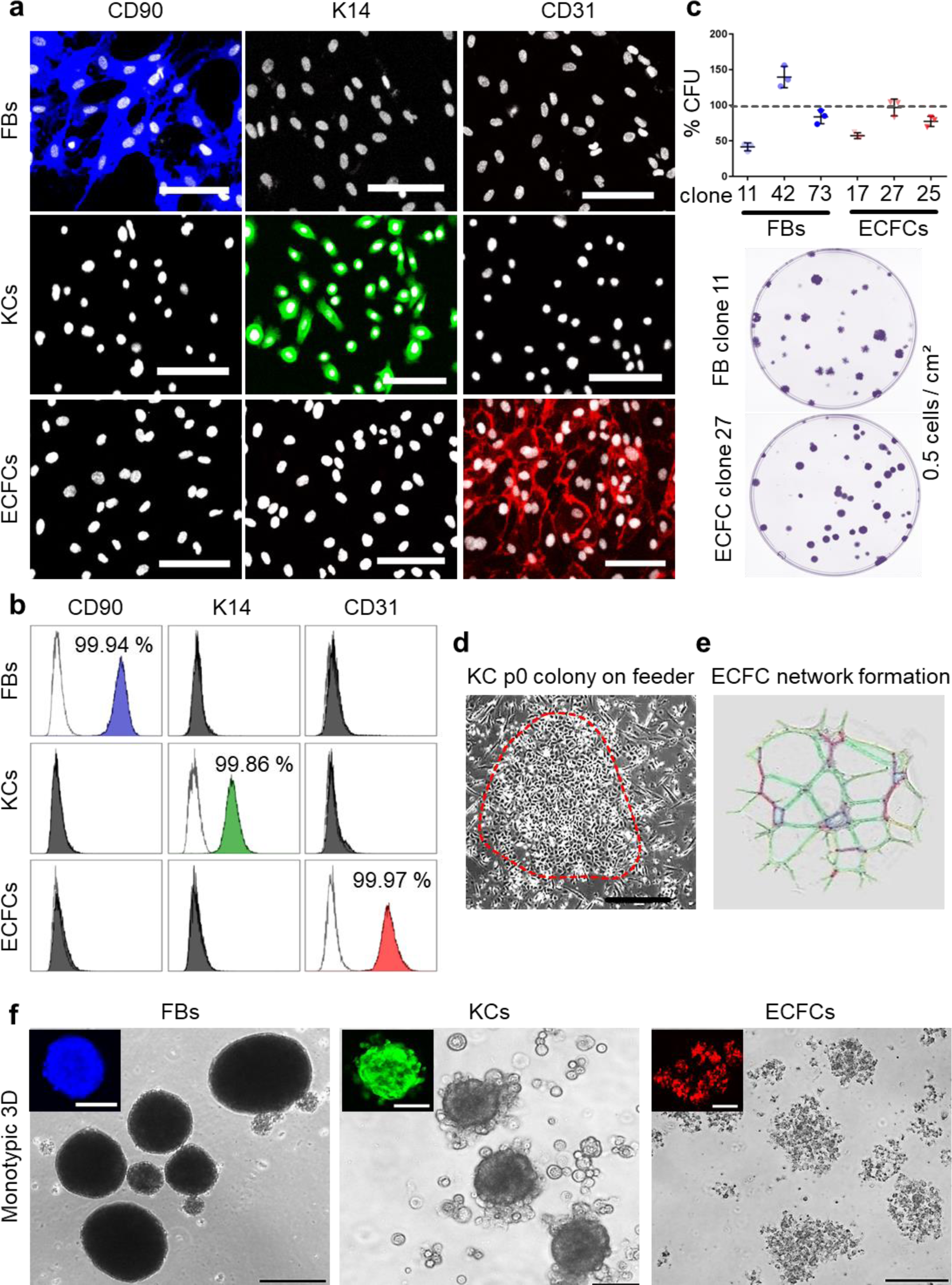
Characterization of purified adult skin and vascular progenitor populations. (**a**) Immune-fluorescence showing CD90 expression of culture-expanded adult skin fibroblasts (FBs, blue), intracellular K14 expression of keratinocytes (KCs, green) and CD31 surface expression of endothelial colony-forming progenitor cells (ECFCs, red; all pseudo-colored). (**b**) Flow cytometry confirmed purity of isolated cells. (**c**) CFU assays showed donor-dependent clonogenicity of FBs and ECFCs (n = 3; mean ± SD). (**d**) KC colony on a feeder layer. (**e**) Vascular network formation after 12 hours on matrigel confirmed angiogenic potential of ECFCs (color-coded for automatic counting). (**f**) Primary skin FBs and KCs, but not ECFCs formed compact monotypic 3D spheroids (FBs = blue-, ECFCs= red-, KCs= green-labeled with nanoparticles). (**a, b, d-f**) Data from one of three independent donors shown. Scale bar = 100 µm.

### Platelet-derived growth factors promote *skin* progenitor cell self-assembly into organoids

Skin organoids were established by proportional seeding of keratinocyte, fibroblast and endothelial single-cell suspensions in appropriate media under ultra-low attachment conditions. Different serum-free and serum-supplemented media supported variable organoid formation efficacies exclusively in the presence of hPL providing platelet-derived growth and regenerative factors, resulting in superior self-organization of dermal fibroblast- and ECFC-containing cores covered by a keratinocyte surface layer (**Fig. 2ab, Fig. S2a**). Using fluorescent nanotracker-labeled cells we detected fibroblast aggregation within <10 hours after seeding, followed by stromal-vascular dermal-like core formation, and keratinocytes superficially settled after >24 hours. Video-imaging visualized consecutive organoid assembly, followed by ECFC arrangement and surface anchorage of KCs (**Fig 2cd, Suppl Video 1**). To challenge mechanistic functionality of skin organoids as in vitro model addressing molecular response signatures, we subjected skin organoids as well as their constituting single cell types to kinase-dependent skin inflammatory IL-17A input signals. Kinome profiling using three different antibody array signatures (n = 11 arrays) constantly showed time-dependent significant deviation of organoid response patterns compared to the respective single cell types. Whereas phosphorylation events occurred as expected for single cells, a surprisingly distinct and reproducible recruitment of signal molecule clusters was found in organoids (**Fig. 2e, Fig. S2bc**).

**Figure 2:**
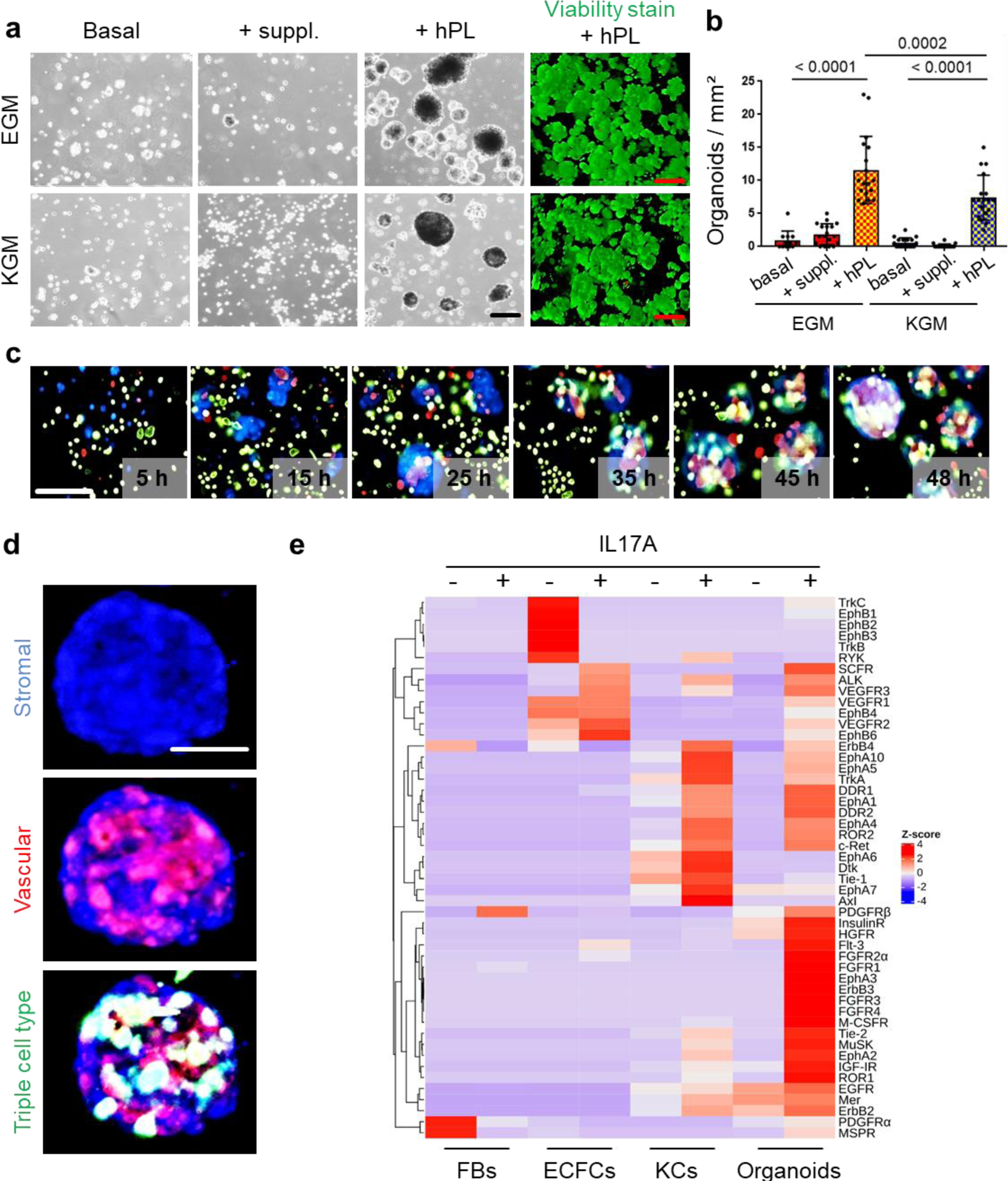
3D organoid organization and functionality. (**a**) Triple cell type organization of fibroblasts, keratinocytes and endothelial cells as evaluated in endothelial growth medium (EGM) and serum-free keratinocyte growth medium (KGM). Light microscopy showed viable (green fluorescence viability dye) organoid formation promoted by platelet-derived growth factors (+hPL), whereas basal media without hPL or with basic supplements (+suppl.) did not support organoid formation. (**b**) Significantly more organoids were formed in the presence of hPL. One-Way-ANOVA and multiple comparison of three areas of two biological and three technical replicates; p values as indicated. (**c, d**) Live cell tracking in the 3D organization process with fluorescent nanoparticles (fibroblasts, blue; endothelia; red; keratinocytes, green). Organoid 3D assembly starting from initial stromal-vascular aggregation and followed by superficial anchorage of adult KCs, indicating well-organized organoids at app. 48 h after cell seeding, repeated in triplicates. Scale bar: 100 µm. (**e**) Proteome profiling of un-starved single FBs, ECFCs, KCs and 4-day assembled organoids after 12 hours stimulation in the absence (HCL control) or presence of IL17A. Z-scores (selected representative analysis).

### Platelet-derived growth factors also revise self-organization of human single-cell suspension transplants in murine full-thickness dermal wounds, creating neo-vascularized human skin

We next determined if self-assembly of primary human skin cells in the absence or presence of ECFCs and platelet-derived factors, respectively, resulted in planar vascularized human skin formation. Xenotransplantation of immune-deficient NOD.Cg-*Prkdc*^*scid*^ *Il2rg*^*tm1Wjl*^/SzJ (NSG) mice was performed after inserting a silicone grafting chamber into full-thickness eight mm diameter skin wounds. Single-cell suspensions of fibroblasts and keratinocytes were either pooled in fetal bovine serum (FBS) as described in a traditional protocol ^33^, or in 10% hPL-supplemented media, in the absence or presence of human ECFCs, immediately prior to grafting the cell suspension onto the murine muscle fascia inside the grafting chamber. Chambers were removed seven days after grafting, allowing the human graft to connect with the murine wound edges (**Fig. S3a**). Skin biopsies taken 14 and 28 days post grafting revealed cell organization into human epidermal and dermal layers (**Fig. 3a-h, Fig. S3b**). Histology confirmed human cell origin and appropriate epidermal and dermal localization in the grafts (vimentin^+^, human dermal cells; keratin 14^+^, human epidermal cells) (**Fig. 3a-d, Fig. S4a**). Proliferating cells were found restricted to basal epidermal keratinocyte layers and scattered dermal cells (**Fig. 3e-h**).

**Figure 3:**
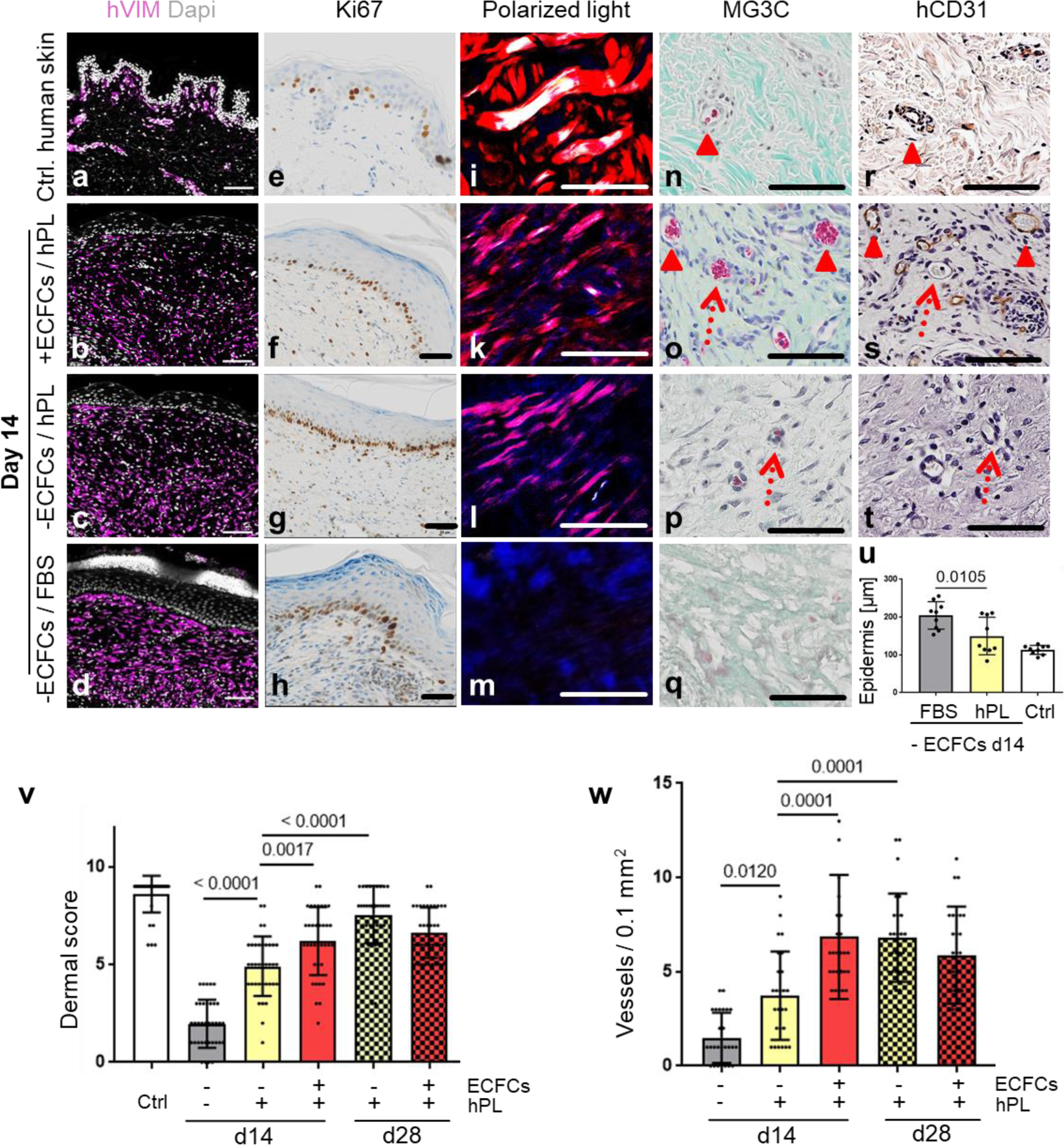
Skin cell self-organization and endothelial cell co-transplantation resulting in properly layered and rapidly vascularized human skin in vivo. Single-cell suspensions (adult keratinocytes, fibroblasts and endothelial cells) re-suspended in 10% hPL- or 10% FBS-supplemented media were transplanted and grafts collected days 14 or 28 (**Fig.S3a**). (**a-t**) Histology of transplants in the absence or presence of ECFCs, supported by hPL or FBS, day 14 post grafting; compared to (**a,e,i,n,r**) healthy human control skin. (**a-d**) Anti-human vimentin (hVIM) confirmed the human origin of the dermis and stratified skin organization; DAPI^+^ nuclei, white. (**e-h**) Anti-Ki67-labelled proliferating cells (brown). (**i-m**) Polarized light-activated collagen fibers exclusively in control skin and hPL-supported, not in FBS-driven transplants **(n-q)**. Masson-Goldner trichrome (MG3C) histochemistry showed vessel enrichment when ECFCs were co-transplanted with fibroblasts/keratinocytes in the presence of hPL (+ECFCs/hPL) compared to transplants without ECFCs (-ECFCs/hPL) showing occasional murine vessel sprouting. Murine erythrocytes (red) confirmed blood circulation inside vessels. **(r-t)** Anti-human CD31 verified human vessel origin in +ECFCs/hPL. (**n-t**) Dotted red arrows = murine vessels. Filled arrowheads = human vessels. (**a-q**) Scale bar 100 µm. One out of three independent grafts per group shown. (**u**) Quantification showing significantly increased epidermal thickness in –ECFCs/FBS compared to –ECFCs/hPL. (**v**) Dermal quality score was significantly increased in hPL-compared to FBS-supported transplants and by ECFC presence day 14 after transplantation. (**w**) Vessel number in grafted cell-derived self-organized human dermis was significantly increased day 14 after ECFC co-transplantation. (**u-w**) One-Way-ANOVA, multiple comparison of three biological and three technical replicates; p values as indicated.

Based on our observation that platelet-derived factors supported progenitor self-assembly into skin organoids in vitro, we next tested different grafting concentrations of hPL (1%, 10%, 100%) and found consistent epidermal-dermal composition with limited erythrocyte extravasation preferentially with 10% hPL-supplemented transplant medium (**Fig. S4b**). Histology quantification showed significant epidermal thickening with signs of hyperkeratosis in FBS-containing transplants reminiscent of inflammatory responses. Epidermis created under the aegis of hPL did not differ significantly in thickness from healthy human abdominal control skin (**Fig. 3u**). Dermal scoring (**Table S1**) was significantly higher in hPL-compared to FBS-containing transplants indicating superior dermal organization with hPL (**Fig. 3v**). Moreover, 28-day skin cell grafts in hPL conditions resulted in more mature dermal architecture as evidenced by significantly increased dermal score compared to 14-day grafts (**Fig. 3v**), comparable to normal skin (**Fig. S4a**). Adding hPL supported normal epidermal development, normal dermal cell distribution, collagen production and murine sprouting angiogenesis compared to FBS, as shown by hematoxylin/eosin and elastica staining (**Fig. S4a**), polarized light stimulation (**Fig 3i-m**) and trichrome histochemistry (**Fig. 3n-q**).

Successful tissue regeneration is strongly dependent on effective vascularization of injured organs, rapidly supporting the tissue with oxygen and nutrients ^8^. Extending previous experience with robust human vasculogenesis induction by combining ECFCs with stromal cells ^11^, co-transplantation of ECFCs significantly increased dermal vessel formation compared to keratinocyte/fibroblast transplants devoid of ECFCs, despite adding hPL in both conditions (**Fig. 3opw**). De-novo formed human vessels, labeled by a human-specific CD31 antibody, co-localized with adjacent in-grown presumably murine vessels lacking CD31 (**Fig. 3s**). Abundance of murine erythrocytes further indicated proper blood circulation and connection of human with murine vasculature (**Fig. 3op**). A summary of transplant groups and key test parameters is given in **Table S2**.

### Sustainable ectodermal lineage specification of hiPSC-KCs augmented by keratinocyte growth factor mRNA transfer

We next aimed to build hiPSC-derived skin cells for testing their self-organization capacity in vivo. Human iPSCs represent a versatile source for differentiated cell propagation for experimental and transplant purposes, particularly in case of limited tissue availability, e.g. after extensive skin burns ^34^. The hiPSC clones were first large scale expanded, generating 1 × 10^9^ cells from less than one million starting cells within 17 days (**Fig. 4a**) while maintaining a pluripotency phenotype (**Fig. 4b**). Three-lineage differentiation was performed in parallel for subsequently testing self-organization of hiPSC-derived fibroblasts (hiPSC-FBs), endothelial cells (hiPSC-ECs) and keratinocytes (hiPSC-KCs). Keratinocytes as ectodermal lineage progeny were differentiated by combining retinoic acid and bone morphogenetic protein as described ^35^, with moderate differentiation (**Fig. S5a**), but limited proliferation upon maturation (**Fig. 4c)**. We thus hypothesized that instant availability of keratinocyte growth factor (KGF, also known as fibroblast growth factor-7, FGF-7) otherwise provided by stromal cells ^21^ could enhance differentiation and proliferation producing sufficient numbers of functional hiPSC-KCs for further testing. The hiPSCs were thus transfected based on a recently established protocol ^36^ with stabilized KGF mRNA during early differentiation. Transfecting hiPSCs with green fluorescent protein (GFP) mRNA as reporter revealed >40% efficiency 24 hours post transfection (**Fig S5b**). Combined transfection of KGF and GFP mRNAs indicated protein expression initiation 0.3 hours after transfection by green fluorescence (**Fig. 4d**).

**Figure 4:**
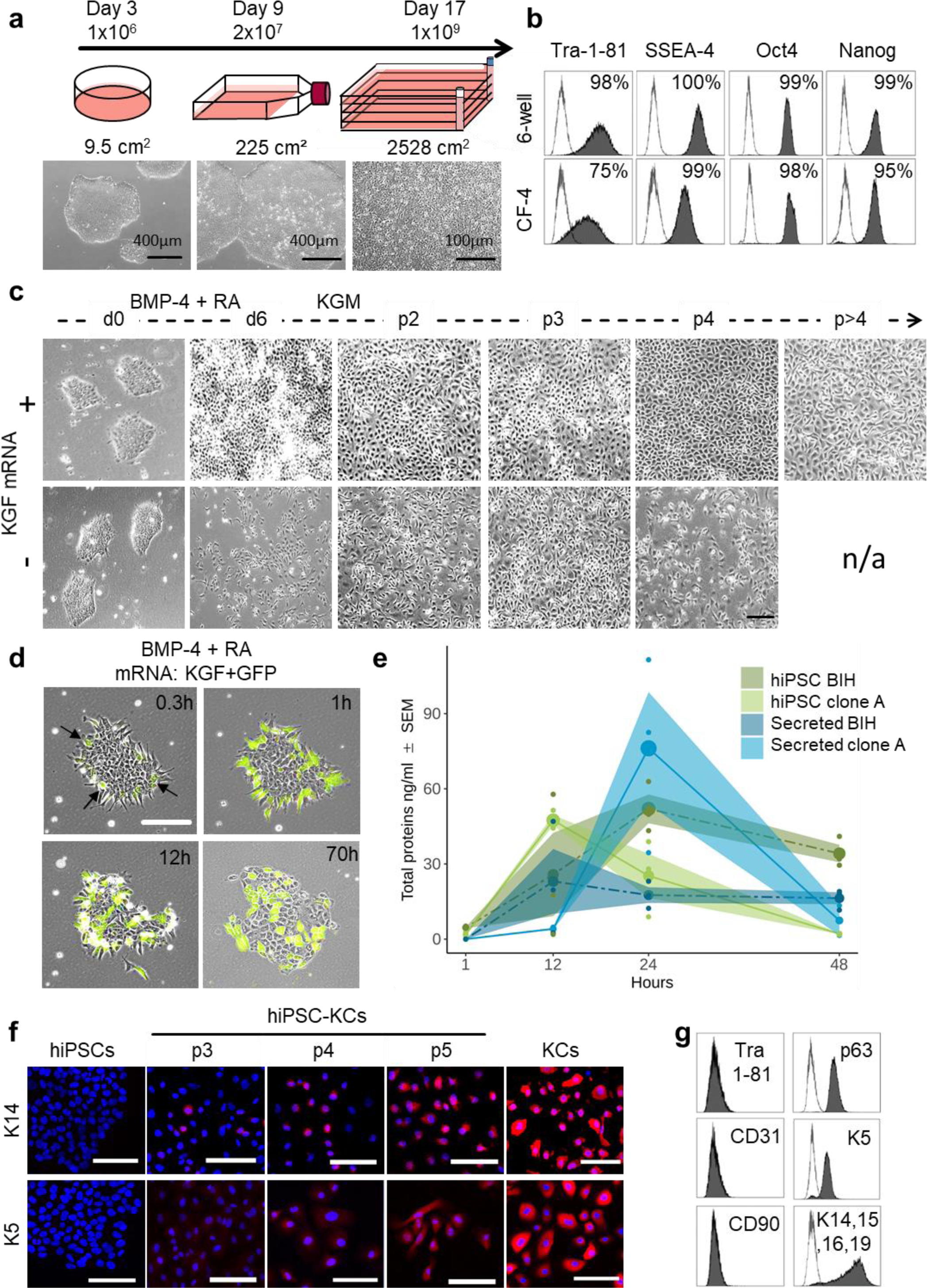
KGF mRNA transfection improved ectodermal cell specification from hiPSCs. **(a)** hiPSC colonies were thawed at day 0 before seeding in a 6-well plate. At day 3 the cells were harvested at 80% confluency (≈1 × 10^6^ cells) and transferred into a T225 flask. At day 9 cells reaching 80% confluency (≈20 × 10^6^ cells) were transferred into a 2,528 cm^2^ cell factory (CF-4). The cells continued to grow clonally and were harvested at day 17 as single cells (≈1 × 10^9^ cells). **(b)** Marker expression (Tra 1-81, SSEA-4, Oct4 and Nanog) of hiPSCs at day 3 and day 17 revealed maintenance of pluripotency upon large scale expansion (one representative sample each, n = 2). (**c**) Morphological changes of hiPSCs during the differentiation into hiPSC-KCs from day 0 to passage >4, in the presence or absence of KGF mRNA during the differentiation phase illustrating limited proliferation potential without KGF. (**d**) Stabilized KGF and GFP mRNA co-transfection indicated protein expression starting 18 minutes after transfection by green fluorescence. (**e**) KGF ELISA confirmed intracellular expression (green) and secretion (blue; n =3, duplicate analysis; individual data points as filled small circles and mean values as large circles connected by lines, ±SEM shaded areas). (**f**) Immunofluorescence staining confirmed rising keratin (K5/K14) expression over time resulting in mature basal progenitor-type hiPSC-KCs. (**g**) Flow cytometry showing Tra 1-81^-^/CD90^-^/CD31^-^, p63^+^, keratin K5^+^/K14^+^/K15^+^/K16^+^/K19^+^ hiPSC-KCs population at passage four (representative staining; n = 2 independent clones).

Intracellular KGF protein expression reached a donor-dependent maximum after 12 - 24 hours. KGF secretion increased with delay (**Fig. 4e**). Appearance of hiPSC clones changed immediately after initiating ectoderm specification into epithelial-like structures (**Fig. 4d, Video S2**). Further maturation of hiPSC-KCs resulted in successive epithelial morphology acquisition, extensive proliferation beyond passage four (**Fig. 4cd**) and stepwise specific keratin (K5, K14) expression (**Fig. 4f**). Flow cytometry confirmed loss of pluripotency markers with assuming a mature hiPSC-KC phenotype comparable with basal KC progenitors (**Fig. 4g, Fig. S6a**).

### Platelet-derived factors promote mesodermal specification of hiPSCs and ameliorate angiogenic potential of hiPSC-derived endothelial cells

For dermal hiPSC-FB generation, mesoderm induction, stromal specification and maturation was performed according to an hPL-based protocol ^37^ (**Fig. S6a, Fig. S7ab**). Mature hiPSC-FBs were rich in clonogenic progenitors (**Fig. S7c**) and could be propagated in 3D (**Fig. S7d**). CD26 (dipeptidyl peptidase-4) expression of mature hiPSC-FBs ranged from 20% to 40%, reminiscent of a neonatal fibroblast phenotype ^38^ (**Fig. S6bc**). For transplant vascularization, endothelial cells (hiPSC-ECs) were generated based on published protocols ^39^ with minor modification (**Fig. S8a**). In situ reporter staining using CD31 antibodies allowed online monitoring hiPSC-EC appearance and maturation in culture, revealing time-dependent increase to up to 55.7% CD31^+^/CD34^+^ hiPSC-ECs until day seven (**Fig. 5ab**). Mean differentiation efficiency was 35.6 ± 3.9% CD31^+^ cells before and 99.7 ± 0.2% CD31^+^ after cell sorting (**Fig. 5c, Fig. S8b**) with a mature Tra 1-81^-^, SSEA^-^, CD90^-^ and CD73^+^, CD105^+^, CD31^+^ EC phenotype (**Fig. 5d, Fig. S6a**) and clonogenic potential (**Fig. 5e**). In this study, we did not test more extensively whether bona fide ECFCs were initiating the colonies ^15,40^ because the focus was on using post-natal and hiPSC-derived endothelia for skin vascularization. Purified hiPSC-ECs formed pronounced vascular networks in the presence but not absence of hPL. To gain further insight into the mechanism of hPL-based angiogenesis support ^11,13^ we purified extracellular vesicles (EVs) from hPL, because recent evidence indicated that EVs can mediate such trophic effects ^41^. The hPL-derived EVs supported angiogenesis of hiPSC-ECs significantly in a dose-dependent manner even in the absence of otherwise essential externally added vascular endothelial growth factor (VEGF). Undifferentiated hiPSCs did not respond to pro-angiogenic stimuli (**Fig. 5fg**).

**Figure 5:**
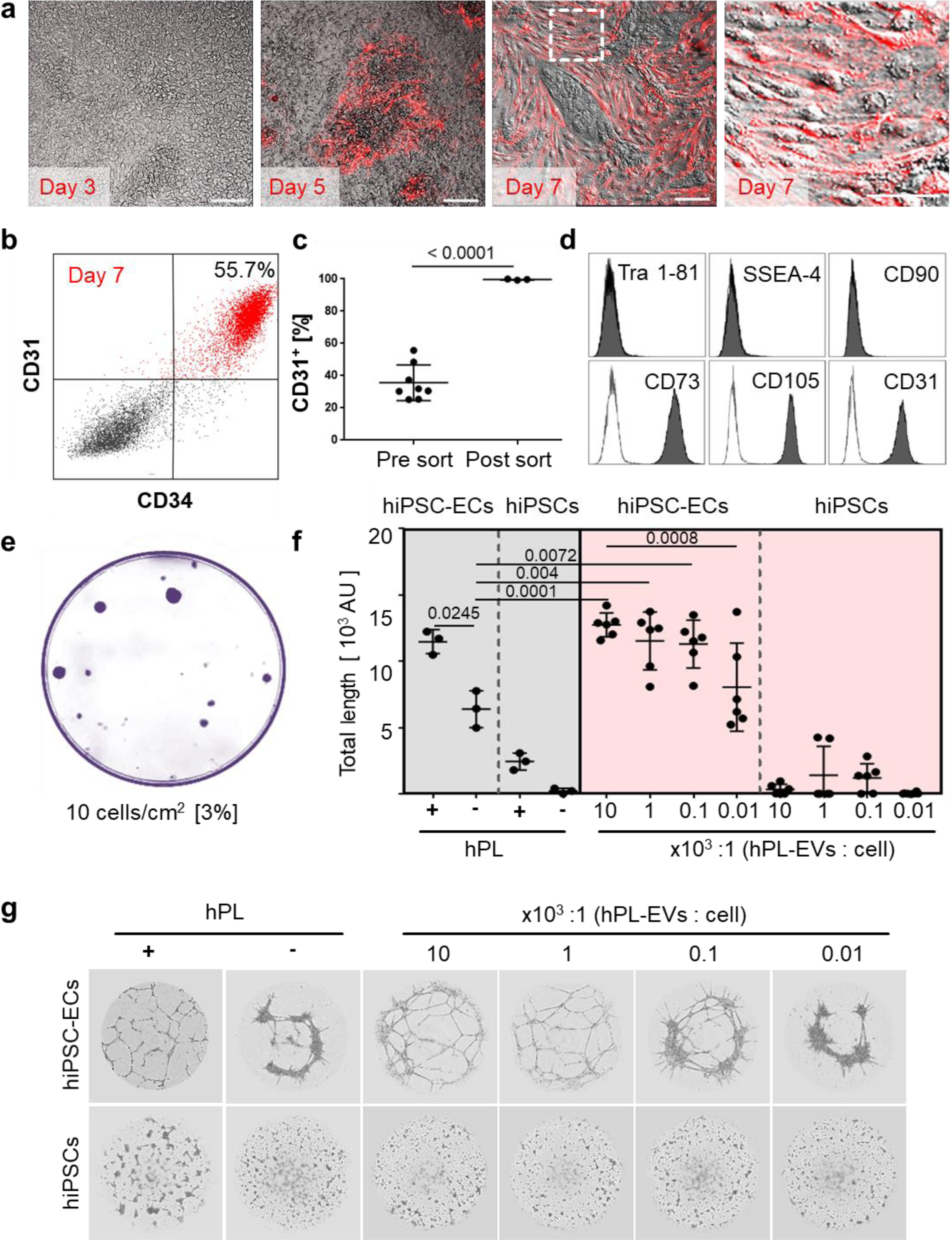
Platelet-derived extracellular vesicles augment angiogenic function of hiPSC-ECs. (**a,b**) *Repetitive in situ* reporter staining of hiPSC-EC cultures during differentiation using an anti-human CD31 antibody highlighting differentiated cells (red), corresponding to (**b**) 55.7% CD31^+^/CD34^+^ hiPSC-ECs in flow cytometry. (**c**) Differentiation efficiency towards CD31^+^ hiPSC-ECs was mean 35.6 ± 3.9% (n=8), and sort-purification led to 99.7 ± 0.2% (n=3). Statistical analysis in GraphPad Prism 7.03, t-test, 3 independent hiPSC clones; p value as indicated. (**d**) Phenotypic analysis showing pure Tra 1-81^-^, SSEA-4^-^, CD90^-^, CD73^+^, CD105^+^, CD31^+^ mature hiPSC-ECs. (**e**) Colony-forming unit assay (CFU) showed 3% clonogenic potential of hiPSC-ECs seeded at a density of 10 cells/cm^2^. Representative example shown, n = 3. (**f**) Vascular-like network formation quantification of total network length showing angiogenesis by differentiated hiPSC-ECs, but not hiPSCs; platelet-derived growth factor (hPL)-dependent; increased network formation with increasing hPL-derived extracellular vesicle (EV) dose. One-Way-ANOVA and multiple comparison of two independent hiPSC donors; p values as indicated. (**g**) Vascular-like networks formed on matrix. Representative examples shown, n = 6.

### In vivo self-assembly of hiPSC-derived single-cell suspension transplants

Ectodermal hiPSC-KCs and mesoderm-derived hiPSC-FBs plus hiPSC-ECs were combined (ratio 2:1:1) as single-cell suspensions in 10% hPL-supplemented transplant medium as established with adult cells and grafted into eight mm diameter full-thickness skin wounds on the back of NSG mice. Analysis was performed 14 days and three months after transplantation (**Fig. 6a-l**). Histology confirmed proper organization into epidermal and dermal layers with proliferating Ki67^+^ hiPSC-derived epidermal cells predominantly in the basal KC layer, still detectable after three months (**Fig. 6ab**). Polarized light microscopy ^42^ highlighted maturing collagen fibers produced by hiPSC-FBs (**Fig. 6c**). In situ labelling of human nuclei using arthrobacter luteus (Alu) repetitive sequences ^43^ confirmed human skin origin (**Fig. 6d**). Histochemistry showed vascularized hiPSC-derived dermis and perfused vasculature after two weeks (**Fig. 6ef**), confirmed by human-specific CD31 staining, filled with murine erythrocytes (**Fig. 6gh**). Contamination of transplants with immature hiPSCs bears teratoma risk. We did not observe teratoma formation within a 3-month observation period in two animals tested, matching requirements previously summarized to minimize tumorigenic risk qualitatively ^44^ Three months after transplantation, macroscopic transplant analysis revealed no aberrant structures at the transplant site (**Table S3**). Histology and Ki67 staining showed skin wound contraction as typically observed in rodents but no signs of malignancy and no highly proliferative structures in the graft (**Fig. 6kl**).

**Figure 6:**
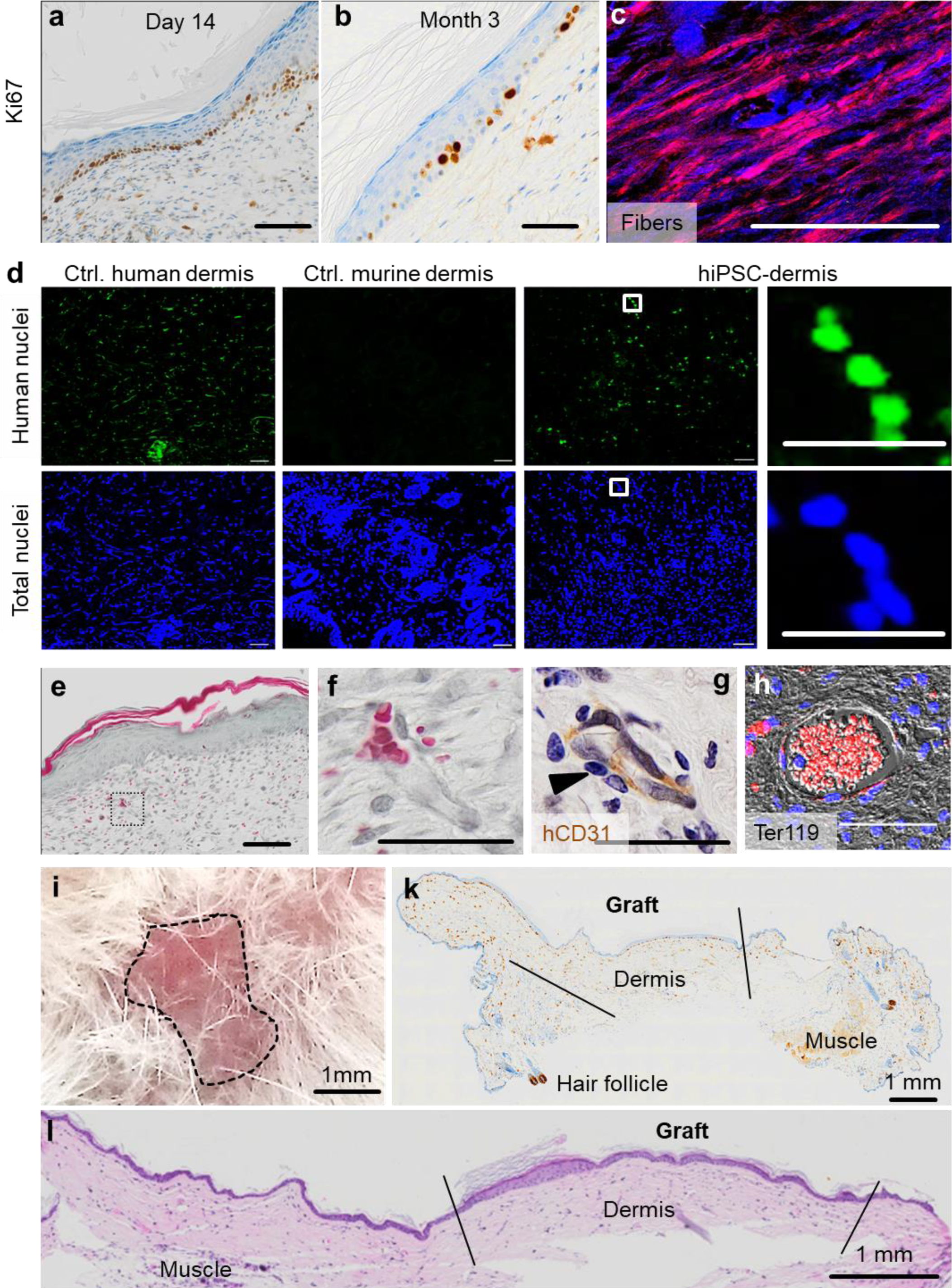
In vivo self-assembly of hiPSC-derived human skin. (**a**) Single-cell suspension transplants of hiPSC-derived cells were analyzed day 14 and (**b**) three month after transplantation. Histology confirmed epidermal-dermal stratification showing proliferating Ki67^+^ basal keratinocytes. (**c**) Polarized light stimulation confirmed maturing collagen fibers day 14. (**d**) In situ hybridization marked human nuclei (Alu sequences, green fluorescence) at day 14. DAPI labelling of total human and murine cell nuclei in blue. (**e, f**) Masson-Goldner trichrome (MG3C) staining showed a vascularized dermis, (**f**) higher magnification, day 14. (**g**) Anti-human CD31 staining labelled hiPSC-ECs along the vessel wall (brown) stabilized by pericytes (blue nuclei in unstained adjacent cells, arrowhead). (**h**) Anti-murine glycophorin-related Ter119 staining confirmed blood supply within vessels (red fluorescence). Scale bar 100 µm. (**i**) Self-assembled hiPSC-derived human skin showed no macroscopic signs of tumor formation three months after grafting (representative picture; n = 2 animals). (**k**) Anti-Ki67, although not human-specific, revealed no aberrant high-proliferative structures three months after grafting (panniculus carnosus, murine muscle; murine hair follicle). (**l**) Hematoxylin/eosin staining confirmed the absence of tumorigenic structures in serial full section scans three months after grafting. (**a-h**) Scale bar=100 µm. n = 2 animals per condition.

## Discussion

Taking advantage of a previously neglected self-assembly potential of progenitor-enriched skin fibroblasts and keratinocytes together with endothelial progenitors, we introduce a novel type of vascularized skin organoids. Self-organization of these spheroid skin organoids was dependent on adding a complex mixture of platelet factors. Translating these xeno-free regenerative conditions into a technically straightforward ‘liquid’ transplantation protocol allowed rapid vascularized full-thickness skin wound healing by adult skin cell suspensions as well as by hiPSC-derived progenitor-enriched skin cells. Significantly increased skin quality was observed upon engraftment in hPL compared to standard animal serum. The transplantation of adult or hiPSC-derived skin cell suspensions opens versatile opportunities for developing novel strategies for skin regeneration. The rapidly assembled skin organoids represent another attractive innovation particularly for individualized and cell type-specific pharmaceutical testing.

The precise selection of contributing mature cell types is an advantage regarding safety of hiPSC-based skin cell transplantation but also represents the major limitation of the current study, because it is lacking hair and skin appendages. In a recent landmark study, complex organoid-like structures containing skin with hair and appendages were generated from hiPSCs with a step-wise protocol. Contaminating cartilage and other extracutaneous fetal tissue may currently limit applicability ^21^. The perfect cell-engineered skin transplant will also contain regionalized appendages, nerves and locotypical hair composition. Combining self-assembly strategies with sophisticated developmental biology and cell sorting tools has the potential to realize such complex endeavours in the near future. Ideally this will result in scar-free regeneration otherwise observed during fetal development ^45^. We may speculate that reduced CD26 expression by hiPSC-FBs, reminiscent of the engrailed^+^ non-scarring fetal fibroblast phenotype ^38^, could indicate reduced scar formation risk after transplantation. Furthermore, availability of human leukocyte antigen (HLA)-matched or universal ß2 microglobulin- or HLA-knockout hiPSC lines will be prerequisite for transplanting immune-competent recipients without the need for immune suppression ^46^.

From a technology point of view, our study offers additional timely application paths. (i) Regarding adult cell-derived spray-on skin ^47^, addition of hPL to transplanted cell suspensions might offer the opportunity to better repair wounds with small amounts of regional cells. (ii) Transfection of stabilized lineage-specific growth factor mRNA ^48^, in our study coding for KGF to improve keratinocyte long-term fitness, may be extended to other cell lineages refining hiPSC-derived cell transplantation strategies. (iii) Two types of reporter assays were tested in our study. Co-transfection of fluorescent protein mRNA (GFP) enabled indirect monitoring of KGF expression as confirmed by protein ELISA. Repetitive addition of fluorescent anti-CD31 antibodies allowed for monitoring endothelial cell maturation. Both strategies are expected to find their way into daily laboratory routine due to their simplicity and reproducibility. (iv) We provided preliminary evidence that spheroid skin organoids created within 3-6 days display an organotypic molecular response not observed in the separated skin cell types. More complex but still precisely defined progenitor cell input into rapidly self-assembled skin organoids will also enable a new level of high throughput pharmaceutical and molecular testing ^26^

## ONLINE METHODS

### Ethics statement and animal welfare

Permissions for human blood and marrow cell collection as well as genetic reprogramming were obtained from the Institutional Review Board Medical University of Graz (protocols EK 19–252, EK 21– 060) and the Ethics Committee of the province of Salzburg (protocol 415-E/1776/4-2014). Samples were collected after written informed consent from healthy volunteers according to the Declaration of Helsinki. Human full-thickness skin was obtained as biological waste material after informed consent as approved by the ethical committee of the region of Salzburg (vote number: 415-E/1990/8-216). Animal trial permission was granted according to Austrian legislation (§26 TVG 2012; animal trial number: BMBWF-66.019/0032-V/3b/2018). Neonatal human dermal fibroblasts were used from different donors (sample IDs 0000480241, 0000480240; CC-2509, Lonza). The adult foreskin fibroblast-derived hiPSC line BIHi001-A was registered and described in detail at https://hpscreg.eu/cell-line/BIHi001-A. Adult bone marrow- and neonatal umbilical cord blood-derived hiPSCs are described elsewhere ^37^. All hiPSC strains used in this study were created by non-integrating Sendai virus-based reprogramming.

### Isolation and 2D propagation of primary cells

Keratinocytes and fibroblasts were isolated from human full-thickness skin from male and female healthy donors with ages ranging from 23 to 56 years (8 donors total). Split-thickness human skin with an area of 5 × 5 cm^2^ was generated using an Acculan 3 Ti-Dermatome (GA670, B. Braun), and incubated in dispase (4942078001, Roche) to separate epidermis from dermis. Epidermal sheets were digested with TrypLE (12605028, Gibco) to release the keratinocytes for subsequent culture in collagen I-coated (C3867, Sigma) plates in epithelial culture medium CntPrime (CnT-PR, CELLnTEC). For in vitro and in vivo experiments, exclusively high proliferative keratinocytes were used. Fibroblasts grew out from the dermis upon incubating punch biopsies in α-MEM (4526, Sigma) supplemented with 10% pooled hPL, 0.6% Dipeptiven (Fresenius Kabi), 0.04% Heparin (BC-L6510, Biochrom) and 1% Penicillin-Streptomycin (P0781, Sigma) (α-MEM/10% hPL) as described ^36^. ECFCs were isolated from umbilical cord blood (3 donors) according to an established protocol ^30^ and propagated in EGM basal medium (CC-3156, Lonza) including all supplements as provided by the manufacturer (EGM-2 medium, CC4176, Lonza) and replacing FBS with hPL. All cells were cultured in humidified nitrogen-controlled incubators at 37°C, 5% O_2_ and 5% CO_2_, and were large scale expanded in four-layered cell factories (CF-4, 140360, Nunc/Thermo). For selected animal experiments, fibroblasts were cultured for comparison in α-MEM/10% FBS, otherwise supplemented as described above, whenever specified in the results section.

### Human iPSCs generation and differentiation

Human iPSCs were reprogrammed from primary stromal cells derived from bone marrow and umbilical cord blood using a non-integrative Sendai viral vector kit (A1378001, Life Technologies) at the Harvard Stem Cell Institute (HSCI) iPSC Core Facility (Cambridge, MA, USA) adapted from an established protocol ^49^ and characterized in a previous project ^37^. In addition, we used hiPSCs from dermal fibroblasts (BIHi001-A), generated and characterized by the Stem Cell Core Facility, Charité, Berlin Institute of Health. Human iPSCs were cultured on Matrigel^®^ (354234, Corning) in mTeSR™ 1 (85850, STEMCELL Technologies) medium. Large-scale expansion to generate a working cell bank for subsequent experiments was performed in four-layered cell factories. For ectoderm induction, 30% confluent dermal and bone marrow derived hiPSC colonies were propagated in keratinocyte serum-free medium (17005042, ThermoFisher) supplemented with 1 µM retinoic acid (R2625, Sigma) and 25 ng/ml bone morphogenetic protein-4 (SRP3016, Sigma) for 6 days performing a media change every second day according to published work ^35^. In addition, the cells were transfected with 1 µg keratinocyte growth factor (KGF) stabilized mRNA (TriLink) at day 0 and day 2 during differentiation. Keratinocyte maturation and expansion were performed in Cnt-Prime epithelial culture medium for approximately 30 days. Mesoderm induction and fibroblast maturation from bone marrow and umbilical cord blood derived hiPSCs, respectively, were performed based on an established protocol ^37^. The maturation phase was extended from passage 4 to passage >8 in α-MEM/10% hPL to obtain mature CD90^+^ hiPSC-FBs. Bone marrow and umbilical cord blood derived hiPSCs were used for endothelial differentiation according to an established protocol with modifications ^39^. In brief, 5 × 10^4^ hiPSCs per cm^2^ were seeded as single cells followed by mesoderm induction using the mesoderm induction medium (05221, STEMCELL Technologies) for 48 hours. Thereafter, the cells were exposed to EGM-2/10% hPL with 260 ng/ml vascular endothelial growth factor (293-VE, R&D Systems) and 2 µM forskolin (F6886, Sigma) for 5 days prior to cell sorting. After sorting, cells were cultured in EGM-2/10% hPL and expanded for characterization, functional assays and transplantation.

### Transfection of in vitro transcribed stabilized KGF mRNA

In order to enhance ectoderm induction and improve long-term growth of hiPSC-KCs, the hiPSCs were transfected simultaneously with eGFP (as a reporter) and stabilized KGF mRNA (TriLink BioTechnologies). The KGF sequence was obtained from https://www.ensembl.org/index.html and produced by using wild type bases. Modification to stabilize the mRNA was done by capping (Cap 1) using CleanCap™ AG / Polyadenylation (120A). Upon DNAse and phosphatase treatment, the mRNA was purified using a silica membrane. Transfection of hiPSC colonies was done using Lipofectamine MessengerMAX transfection reagent (LMRNA003, ThermoFisher) as described ^36^.

KGF protein expression was measured in a time course after KGF mRNA transfection as specified in the results section. For each time point, supernatant was collected and cell protein lysate was isolated using a RIPA Lysis buffer (R0278, Sigma) including a protein inhibitor (87786, Life Technologies). ELISA measurement was performed according to the manufacturer’s protocol (DuoSet ELISA, DY251, R&D Systems) using an OD of 450nm. The optical density values of undiluted and diluted (1:10) samples were calculated using a linear standard curve.

### Flow cytometry and cell sorting

The phenotype of primary cells, hiPSCs and hiPSC-derived cells was identified by flow cytometry using various surface and intracellular antibodies (**see Table S4**). Flow cytometry was performed as previously described ^11,27,50^ with a five-laser BD LSR-Fortessa™ (BD Biosciences), BD FACSDiva Software 8.0.1 Firmware version 1.4 and Kaluza analysis software version 1.3.14026.13330 (Beckman Coulter). In order to obtain pure CD31^+^ endothelial cell populations after seven days of differentiation, cells were labelled with anti-human CD31-eFluor 450 (clone WM59, eBioscience) and subjected to sterile fluorescence-activated cell sorting, using a BD FACSAria™ III instrument (BD Biosciences) and BD FACSDiva Software 8.0.1 Firmware version 1.3.

### Immunofluorescence staining of differentiated cells in situ

For cell characterization, 1 × 10^5^ primary hiPSCs or hiPSC-derived cells were seeded in 12-well plates, grown to 50–70% confluence and fixed with 4% formaldehyde for 10 minutes at room temperature. Cells were washed in PBS and permeabilized in citrate buffer and blocked with 1x Dako wash buffer/10% FBS (S300685-2, Agilent). Anti-human CD90-PE (40 ng/µl, clone 5E10, BD Biosciences), anti-human CD31-PE (12.5 ng/µl, clone WM59, BD Biosciences), anti-human Cytokeratin 14 (4 ng/µl, clone LL001, Santa Cruz), anti-human Cytokeratin 5 (20 ng/µl, clone 2C2, Thermo Fischer) primary antibodies and appropriately titrated isotype controls were applied overnight at 4°C. As secondary antibody, a goat anti-mouse PE (40 ng/µl, BD Biosciences) was applied for 1 hour at room temperature. Cell nuclei were stained with 4′,6-Diamidin-2-phenylindol (DAPI, 1:1000, D1306, Molecular Probes) at room temperature for 10 minutes. In situ reporter staining during endothelial cell differentiation was performed by adding 125 ng/ml of anti-human CD31-PE (clone WM59, BD Biosciences) antibody in basal medium and incubating for 30 min at 37°C. After washing the cells with basal medium, EGM-2 / 10% hPL was added back before reporter staining was analyzed by fluorescence imaging.

### Spheroid and organoid formation 3D

Fibroblasts, keratinocytes and endothelial cells were labelled with cell tracker fluorescent probes (C2110 excitation (ex) / emission (em) 353/466 nm, C7025 ex/em 492/517 nm, C34552 ex/em 577/602 nm; ThermoFisher) according to the producer’s recommended protocol. The spheroid formation capacity of each cell type separately was evaluated by seeding 4 × 10^5^ cells in 6-well ultra-low attachment culture plates (CLS3471; Corning) in the respective propagation media. For skin organoid formation, a single-cell suspension consisting of 1.25 × 10^5^ fibroblasts, 1.25 × 10^5^ endothelial cells and 2.5 × 10^5^ keratinocytes was prepared based on preliminary titration and seeded in various media conditions: (i) keratinocyte serum-free medium basal (17005075, Gibco), (ii) keratinocyte serum-free medium basal + supplements (EGF + BPE), (iii) keratinocyte serum-free medium basal + supplements + 10% hPL, (iv) endothelial cell basal medium (EBM, CC-3156, Lonza), (v) endothelial growth medium (EGM) = EBM + supplements (EGF, IGF, hFGF, VEGF, hydrocortisone, ascorbic acid; CC4176, Lonza), (vi) EGM incl. supplements + 10% hPL. Five days after cell seeding, areas of 1 mm^2^ were counted to determine organoid numbers. Life cell imaging was performed for the first 48 hours; microscopic analysis was done 5 days after seeding (see image acquisition).

### Proteome profiler arrays

Protein isolation and proteome profiler arrays (Human Phospho Kinase array Kit, ARY003C, Human Phospho-RTK Kit, ARY001B; Proteome Profiler Human NFκB Pathway Array, ARY029; all R&D Systems) were performed according to the manufacturer’s protocol comparing stimulation vs. solvent treatment. In brief, for the human Phospho Kinase array, human skin equivalents, skin organoids, fibroblasts, keratinocytes and endothelial cells were starved in serum-free medium for 2 hours before stimulation with 100 ng/ml recombinant human Interleukin 17A (317-ILB-050, R&D Systems) dissolved in 4 mM HCL. For the human Phospho-RTK Kit and the human NFκB Pathway Array non-starved cells and organoids were used. Protein content was measured using an DC Protein Assay Kit II (500-0112, BioRad) according to the manufacturers protocol. Membrane dots were visualized and quantified using a chemidoc system and image lab 6.0.1 software (all Bio-Rad).

### Single-cell suspension grafting

For in vivo grafting, immune-deficient NOD.Cg-Prkdcscid Il2rgtm1WjI/SzJ mice (614NSG, Charles River) were used at an age of 9, 12, 17, 19, 20, 21 and 24 weeks. Full-thickness skin wounds covering app. 2.2% of the body surface area were induced by 8 mm punch biopsy (48801, PFM medical) in the back skin. A silicone grafting chamber including three pores of 2 mm diameter for transplantation and subsequent air contact (50.2 mm^2^ silicone chamber, self-produced; see **Fig. 3a**) was placed on the muscle fascia into the wound. For initial experiments, single-cell suspension transplants consisting of 6 × 10^6^ keratinocytes and 6 × 10^6^ fibroblasts diluted in 200 µl α-MEM/10% FBS (total volume 400 µl) were transplanted as adapted from a published protocol ^33^. In order to enhance the grafting strategy, FBS was replaced by 1%, 10% or 100% hPL and 3 × 10^6^ endothelial cells were co-transplanted as indicated together with to 3 × 10^6^ fibroblasts and 6 × 10^6^ keratinocytes. In addition, hiPSC-derived single-cell suspensions consisting of 6 × 10^6^ hiPSC-KCs, 3 × 10^6^ hiPSC-FBs and 3 × 10^6^ hiPSC-ECs, diluted in 200 µl α-MEM/10% hPL, were grafted. The chambers were removed seven days after transplantation. Skin biopsies were taken at 14 days, 28 days and 3 months after initial grafting, fixed in 4% formaldehyde and prepared for histology.

### Histology of skin sections

Paraffin-embedded skin samples were cut into 4 µm sections for immunofluorescence, histochemistry and immune-histochemistry. For evaluating the human origin of skin graft cells, a mouse monoclonal anti-human vimentin antibody (1:100, clone V9, M072501-2, Dako) followed by a goat anti-mouse PE (40 ng/µl, BD Biosciences) was used. Murine erythrocytes were identified using a rat monoclonal anti-murine Ter119-PE antibody (40 ng/µl, clone TER-119, BD Pharmingen). For human vessel staining, a mouse monoclonal anti-human CD31 antibody (1:100, clone JC70A, Dako) detected by the Avidin-Biotin Complex Kit (ABC System, SP-2002, Vector Laboratories) and developed by diaminobencidine staining (DAB plus Chromogen Solution, K3468, Dako) was used. Basal keratinocytes were detected by a mouse monoclonal anti-cytokeratin 14 antibody (4 ng/µl, clone LL001, Santa Cruz), followed by ABC and DAB detection and development. A rabbit monoclonal anti-Ki67 antibody (clone 30-9, 790-4286, Roche) was used to detect the proliferative cells. Hematoxylin and eosin (H&E) staining was done in a linear slide stainer (Leica ST4040) using Mayer’s Hemalaun (1.09249.2500, Merck) and Eosin Y (1.15935.0100, Merck). For Masson-Goldner trichrome and elastica staining, kits were used according to the manufacturer’s recommendations (12043, 14604, Morhisto). Collagen fibers in HE stained sections were stimulated with polarized light, using an Olympus™ Rotatable Analyzer (U-AN360-3) filter. Human cell nuclei were labeled via in situ hybridization as previously published ^43^.

### Image Acquisition

Life cell imaging and video recording was performed on an Eclipse Ti inverted microscope (Nikon) with a customized live cell incubation system (Oko lab). Images were taken every 15 minutes for 48 hours. Image analysis was done using NIS elements imaging software AR 4.30.02 (Nikon). Confocal microscopy was performed using laser scanning microscopes Axio Observer Z1 attached to LSM700 (Carl Zeiss) Light microscopic cell culture pictures were generated with an EVOS XL microscope (Thermo Fisher). Total slides were scanned automatically in 40x magnification using the VS-120-L Olympus slide scanner 100-W system and processed using the Olympus VS-ASW-L100 program. Figures were prepared using Microsoft Power Point 2016 and GNU Imaging Program (gimp-2.8).

### Clonogenicity, EV concentration and Network formation assay

The clonogenic potential of fibroblasts, endothelial cells and hiPSC-derived cells was assessed according to published protocols ^50,51^. The purification of extracellular vesicles from hPL supporting the angiogenic potential of ECs and hiPSC-ECs as well as the network formation assay was previously established and performed according to a published protocol ^41^.

### Quantification, dermal scoring, statistical analysis

Vessels were quantified in dermal areas of 0.1 mm^2^ followed by counting the total number of vessels (murine + human vessels) in H&E stained sections using the Olympus VS-ASM-L100 program. The epidermal thickness was measured using the same program. The dermal score was evaluated to grade the dermal quality of the transplants, based on the density of fibroblasts, produced matrix, vessels, hemorrhage and collagen fibers produced. Areas of 0.1 mm^2^ were randomly selected and scored from 0 to 9 points (see **Table S3**). Per group, three to five biological and at three technical replicates were included. Statistical analysis was performed using One-Way ANOVA analysis and multiple comparison in GraphPad Prism version 7.03. For proteome analysis background signals were subtracted and normalized compared to the reference spots locating on the arrays. Probes showing a normalization signal lower than 0.06 in the untreated samples were removed from analysis. Values were scaled using Z-score row scores and mapped on heatmaps using the R package “ComplexHeatmap” ^52^.

## Author contribution

*Conception and design*: PP, DS; *Development of methodology*: PP, LK, MW, BV, CS, DS; *Acquisition of data*: PP, LK, MW, AH, KM, BV; *Analysis and interpretation of data*: PP, LK, MW, BV, RP, DS; *Writing of the manuscript:* PP, DS; *Review and/or revision of the manuscript*: HS, AS, KS, DS; *Administrative, technical, or material support*: CS, ER, HS, KS; *Study supervision*: DS.

## Declarations

AS is an employe of Accanis biotech. The authors do not declare any conflict of interest.

## Data availability statement

All data supporting the findings of the present study are available from the corresponding author on reasonable request.

## Acknowledgements

We like to thank the sponsors of our research: European Union’s Horizon 2020 research and innovation program (grant agreement no. 733006 to DS and no. 731377 to LK and KS), Land Salzburg IWB/EFRE 2014 - 2020 P1812596 and WISS 2025 20102-F1900731-KZP EV-TT (to DS and KS). We are grateful to Sigrid Kahr, Alina Sonderegger and Margit Wieneroiter for excellent technical assistance. We wish to particularly thank Jeroen Bremer (Department of Genetics, University Medical Center Groningen, The Netherlands) for providing templates for silicon chamber production.

## Supplementary Figures

**Figure S1:**
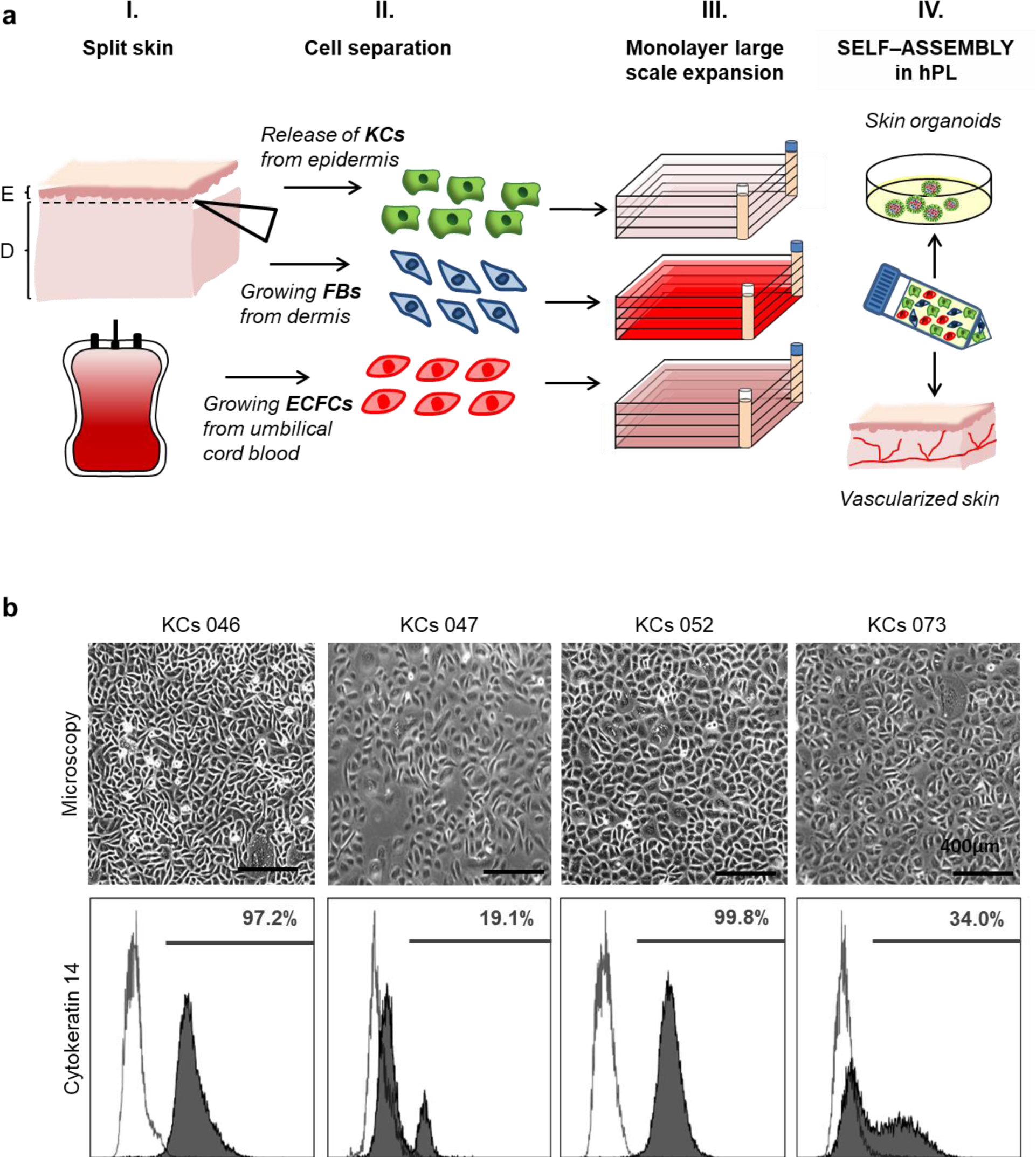
Adult skin cell isolation, characterization and regeneration strategy. (**a**) Illustrative summary of adult keratinocyte (KC), fibroblast (FB) and endothelial colony-forming progenitor cell (ECFC) propagation for testing self-assembly in vitro and in vivo. KCs and FBs were isolated from split-thickness skin explants. ECFCs were isolated from umbilical cord blood (I. + II). All cell types were propagated as monolayers under animal-serum free conditions generating app. 1 × 10^8^ cells per cell factory (III.), before generating single-cell suspensions in human platelet lysate (hPL)-supplemented media promoting cell self-assembly into human skin organoids and vascularized human skin (IV.). (**b**) Morphology and cytokeratin 14 expression of four randomly selected KC preparations illustrating donor variability (Scale bar 400 µm). For cell transplantation, >99% pure cytokeratin 14^+^ keratinocytes were used.

**Figure S2:**
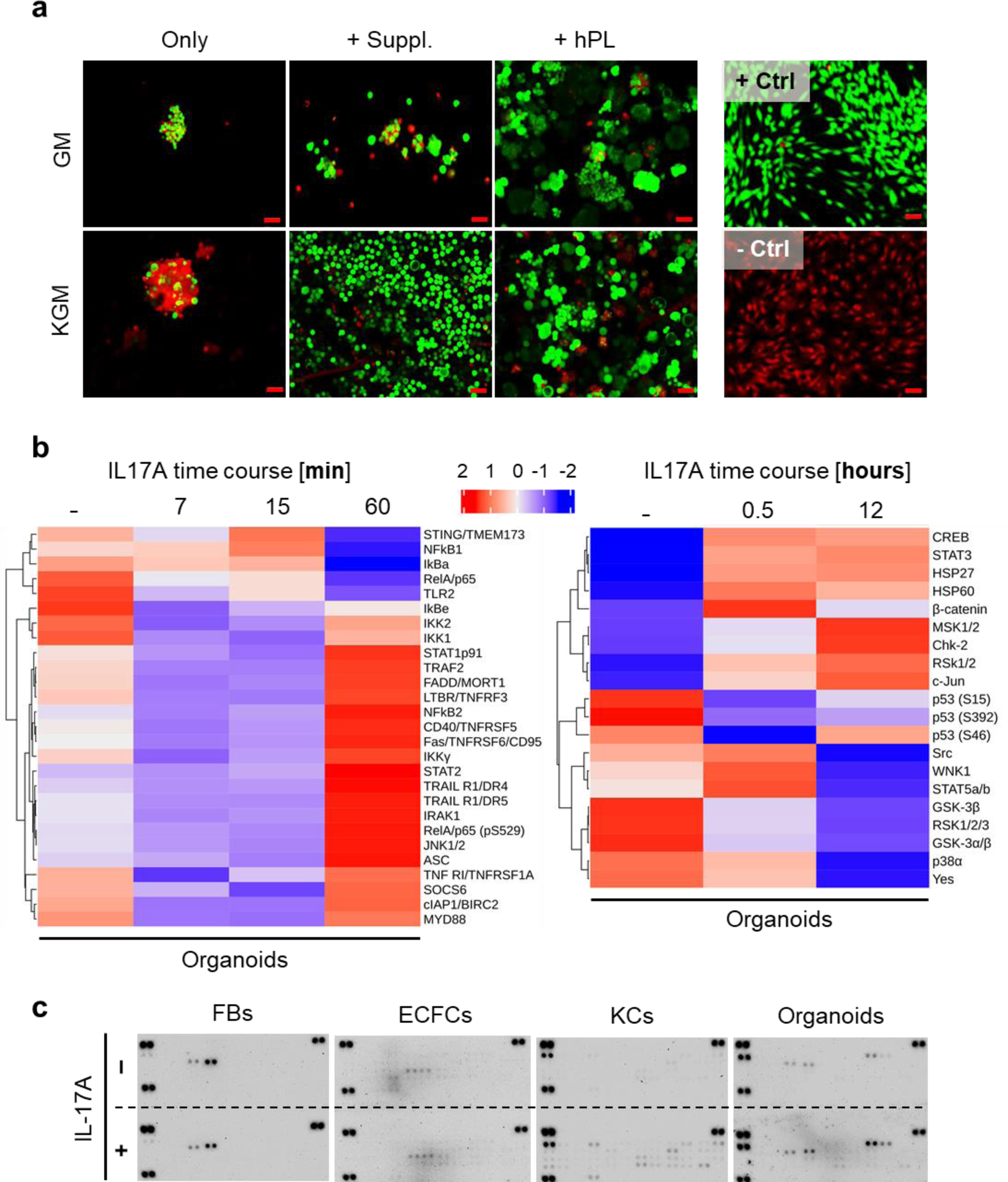
Viability and molecular response of organoids vs. separated cells. (**a**) Life/dead staining (green = viable cells, red = dead cells) showed that viable spheroids are exclusively formed in the presence of human platelet lysate (+ hPL; +ctrl, healthy fibroblasts; –ctrl, ethanol treated fibroblasts; scale bar = 50 µm). (**b**) Proteome profiling early (left) and delayed (right) time course heatmap of organoids in response to IL-17A. Z-score values. (**c**) Representative comparative antibody arrays of single FBs, ECFCs, KCs and 4-day assembled organoids after 12 hour stimulation in the absence (HCl control) or presence of IL17A.

**Figure S3:**
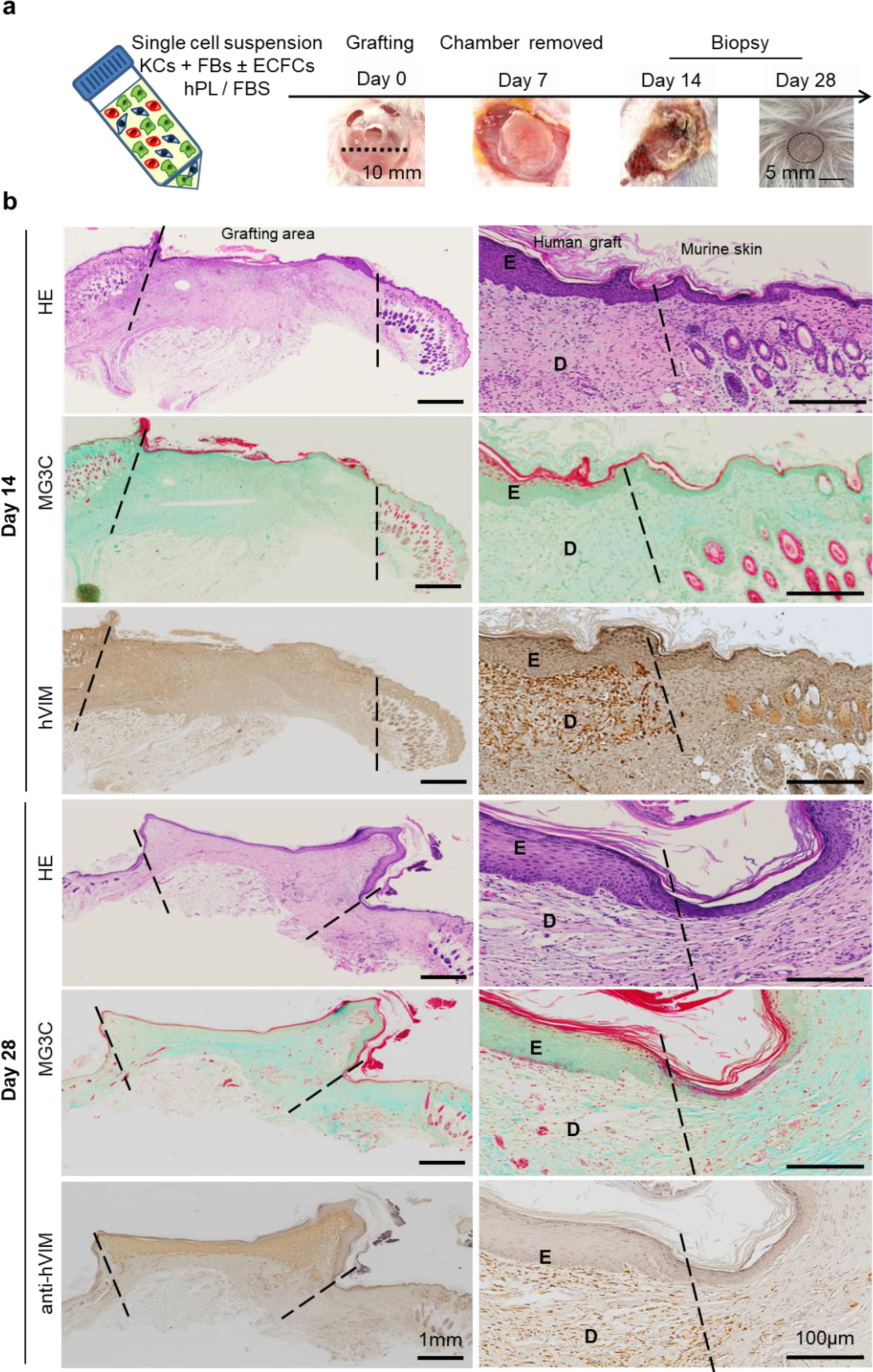
Single-cell suspension transplantation produced layered human skin. (**a**) Grafting procedure illustration: single-cell suspensions (KCs + FBs ± ECFCs) in 10% hPL- or FBS-supplemented medium were transplanted in a silicone chamber inserted in an 8 mm full-thickness wound of NSG mice. Chambers were removed at day 7 and grafts analyzed days 14 or 28. (**b**) Full scan of skin sections 14 (n = 15 animals) and 28 days (n = 12 animals) after transplantation: Hematoxylin (HE), Masson Goldner trichrome (MG3C) and anti-human vimentin (hVIM) staining showing the grafted area surrounded by murine skin (marked by hatched lines). Representative stainings shown.

**Figure S4:**
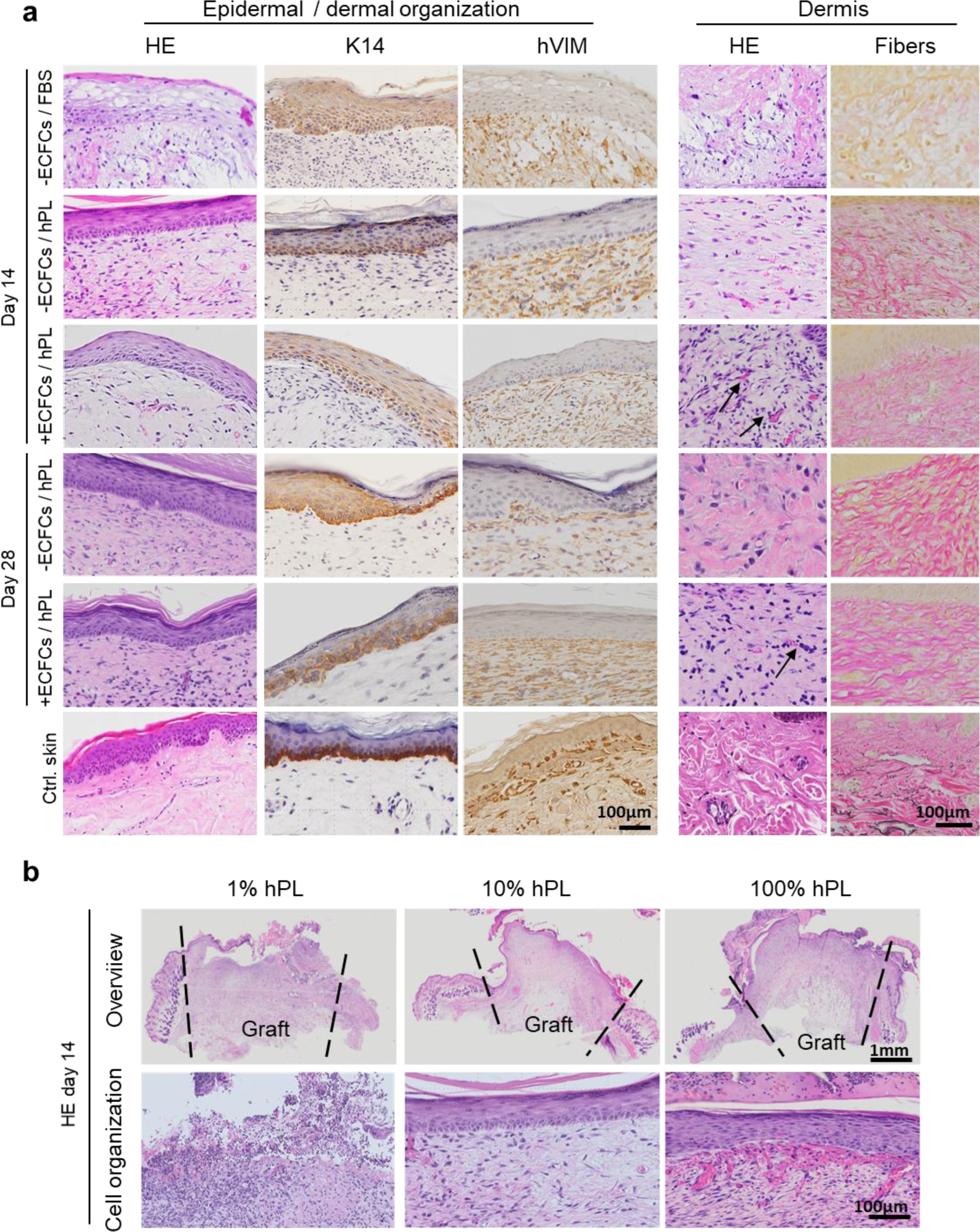
Histology of transplanted human skin 14 and 28 days after engraftment. (**a**) Layered epidermal-dermal cell organization visualized in hematoxylin/eosin staining (HE) and human vimentin (hVIM) dermal cell origin confirmed in all transplant conditions (as identified left of the pictures). Anti-cytokeratin K14 staining showed keratinocytes located predominantly in the basal epidermal skin layer day 28 after grafting. Dermal tissue composition in HE staining and fiber stain (black arrows, vessels; scale bar, 100 µm; representative staining of one out of three independent grafts per group shown. (**b**) Cell grafts consisting of fibroblasts (FBs) and keratinocytes (KCs) in α-MEM/1% or 10% or 100% human platelet lysate (hPL) were transplanted and analyzed after 14 days. Full section scans of HE stained grafts (overview) showed the human graft surrounded by the murine skin (marked by hatched lines). Higher magnification revealed regular skin cell organization at 10% hPL and bleeding in 100% hPL grafts. Data of one pilot experiment shown (n = 3 animals).

**Figure S5:**
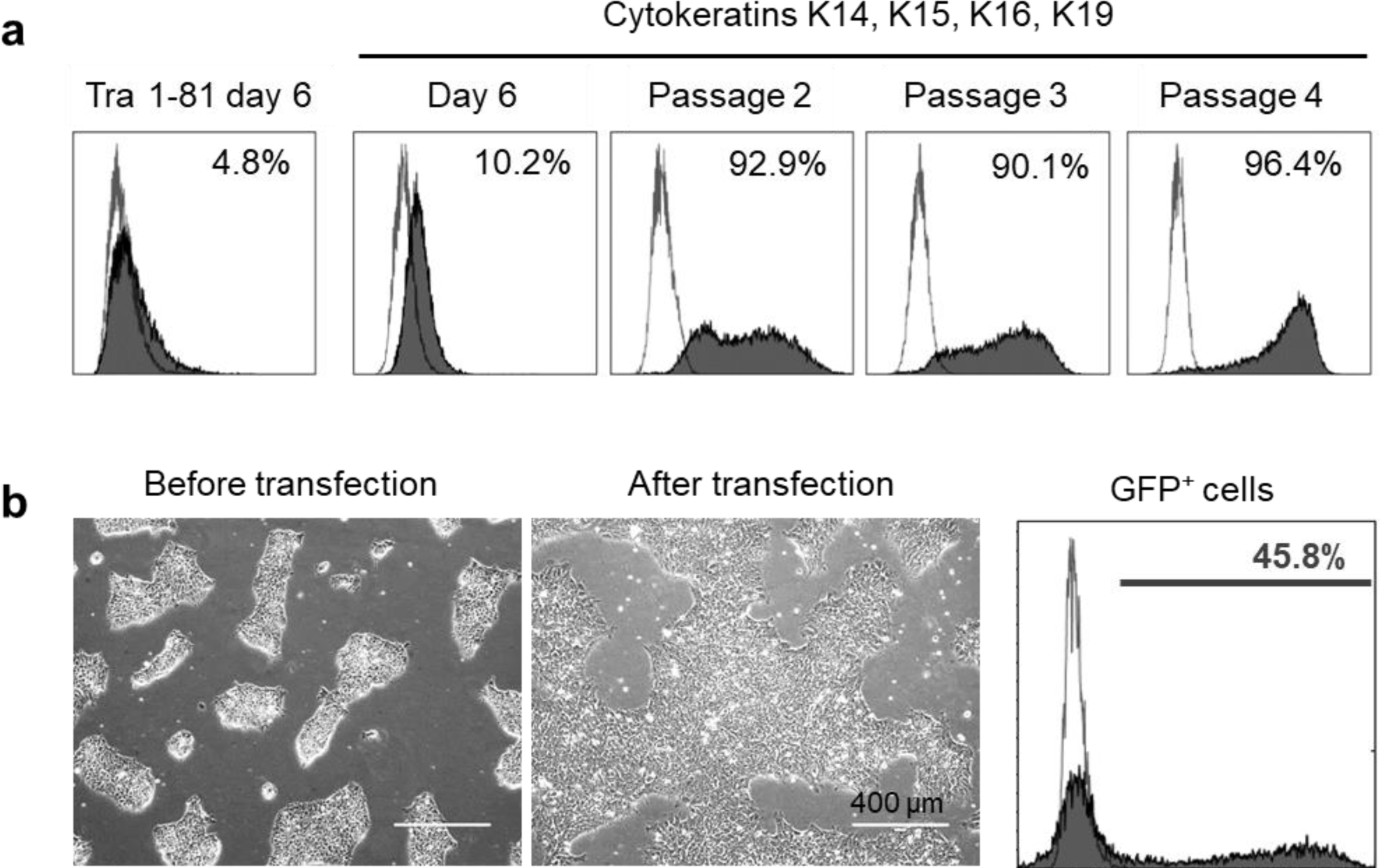
Keratinocyte differentiation from hiPCSs without KGF. (**a**) Reproducing the keratinocyte differentiation protocol containing BMP-4 and RA during the differentiation phase ^35^. Flow cytometry revealed a reduction of pluripotency marker Tra-1-81 until day 6 of differentiation and a stepwise increase in keratinocyte marker expression during maturation from day 6 to passage 4. (**b**) Stabilized mRNA transfection of hiPSCs: Transient transfection of hiPSC clones using a GFP mRNA showed normal cell growth one day after transfection, resulting in 45.8% efficiency 24 hours after transfection. (**a+b**) Data of one pilot experiment shown.

**Figure S6:**
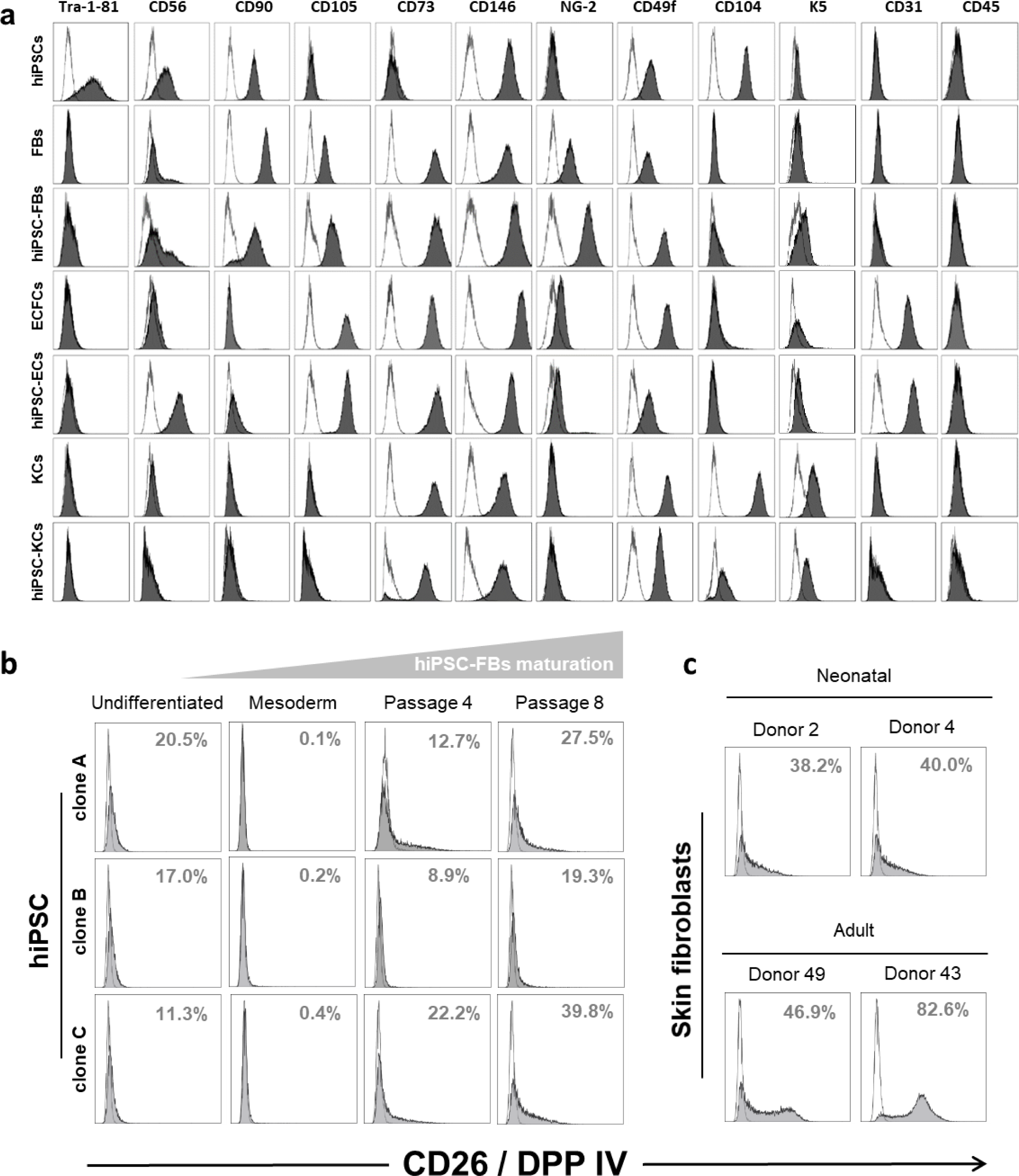
Phenotypic analysis of hiPSC-derived and adult cells and monitoring of hiPSC-derived, neonatal and adult fibroblast’s CD26 expression. (**a**) Flow cytometry showed comparable phenotypes of hiPSC-derived cells and their corresponding adult cell types (n = 1 to 3). (**b**) Increase of CD26 expression during hiPSC-FB maturation, confirming a consecutive transition from a hypothetically regenerative (CD26^-^) to scarring (CD26^+^) fibroblast phenotype during cell maturation. (**c**) CD26 expression levels of neonatal and adult fibroblasts compared to hiPSC-FBs. Two out of three donors shown.

**Figure S7:**
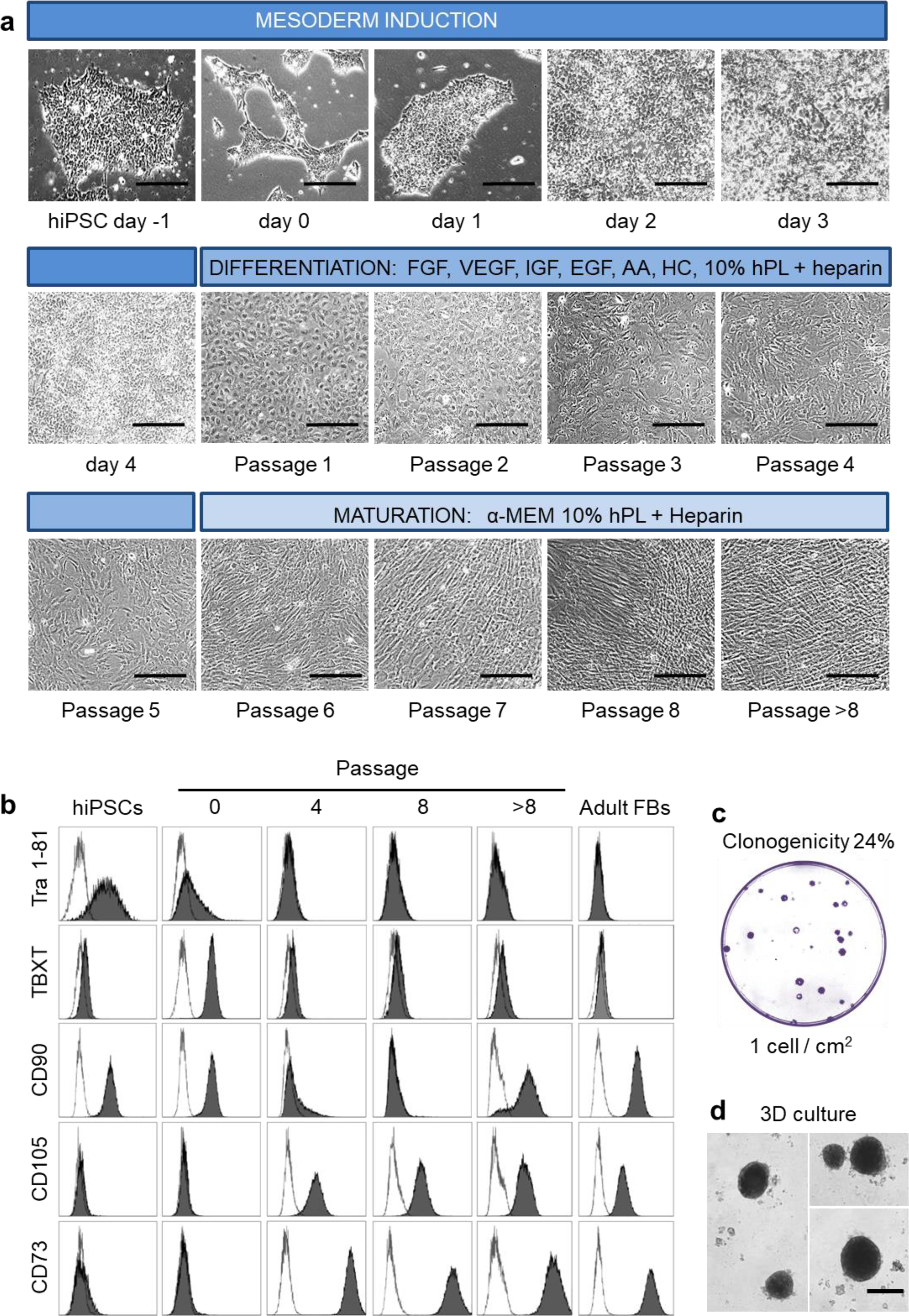
Mesoderm induction, differentiation and maturation of fibroblasts from hiPSCs. (**a**) Morphological changes of hiPSCs during four days of mesoderm induction, followed by five passages of early fibroblast (FB) differentiation and 4-6 passages of FB maturation. **(b)** Phenotypic analysis showed the stepwise differentiation and maturation of hiPSC-FBs differentiation, resulting in Tra 1-81^-^, TBXT/brachyury^-^, CD90^+^, CD73^+^ and CD105^+^ mature hiPSC-FBs assuming a phenotype otherwise comparable to adult FBs. (**c**) Colony-forming unit (CFU) assay showed clonogenic potential of hiPSC-FBs. (**d**) 3D culture of hiPSC-FBs (representative pictures shown). (**a-d**) Representative data from two independent hiPSC clones.

**Figure S8:**
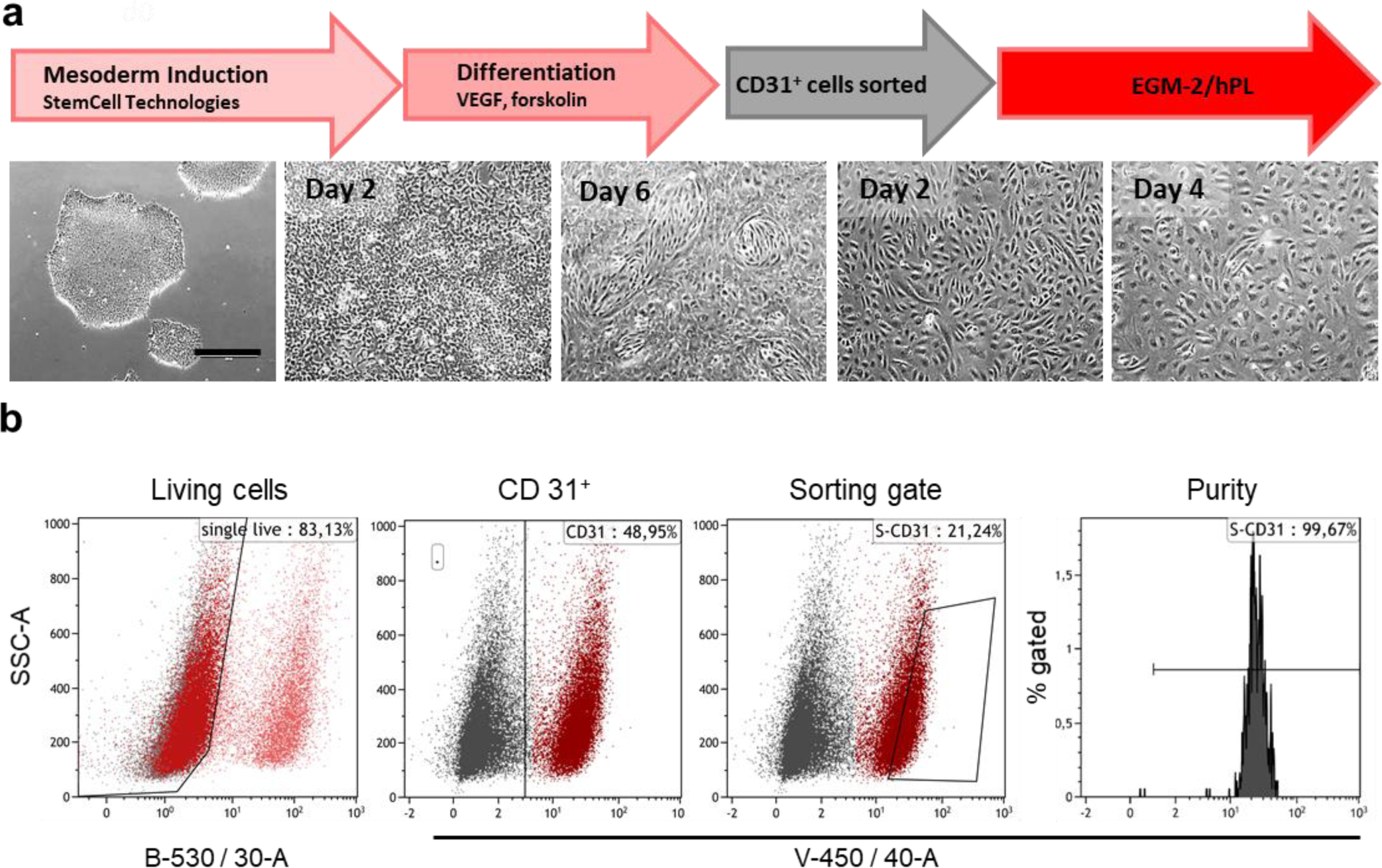
Mesoderm induction, differentiation and maturation of endothelial cells from hiPSCs. (**a**) Morphological changes of hiPSCs during two days of mesoderm induction, followed by 4 days of endothelial differentiation. Day six CD31^+^ cells were purified by fluorescence-activated cell sorting before continuation of endothelial cell maturation. (**b**) Sorting strategy: Viable single cells in this randomly selected sample contained 48.9% CD31^+^ cells in a bulk population at day 6 during differentiation. The sorting gate included 22.8% CD31^+^ cells, resulting a sort purity of 99.5%. (**a+b**) Representative data from one out of three independent hiPSC clones shown.

**Table S1:** Parameters for evaluating the dermal score.

|  |  |
| --- | --- |
| **A) Vascularization and hemorrhage** |  |
| No blood vessels / no hemorrhage | 0 |
| Strong hemorrhage (with or without blood vessels) | 1 |
| Blood vessels / low to moderate hemorrhage | 2 |
| Blood vessels / no hemorrhage | 3 |
| **B) Cell density / ground substance produced** |  |
| High cell density | 0 |
| Moderate cell density / no ground substance | 1 |
| Moderate cell density / moderate ground substance | 2 |
| Low cell density / distinct ground substance | 3 |
| **C) Thickness and density of fibers** |  |
| No fibers | 0 |
| Thin or thick fiber / high fiber density | 1 |
| Thin fiber / moderate to low fiber density | 2 |
| Thick fiber / moderate to low fiber density | 3 |

**Table S2:** Summary of transplant groups and key parameters tested. Absolut numbers or mean ± SD given.

| Transplant group | Cells | FBs+KCs | FBs+KCs | FBs+KCs+<br>ECFCs | FBs+KCs | FBs+KCs+<br>ECFCs | Human<br>skin |
| --- | --- | --- | --- | --- | --- | --- | --- |
|  | Medium | 10% FBS | 10% hPL | 10% hPL | 10% hPL | 10% hPL | - |
|  | Time point | 14 days | 14 days | 14 days | 28 days | 28 days | adult |
| Transplant quality | Human<br>origin | 4/4 | 3/4 | 4/4 | 3/3 | 3/4 | - |
|  | Cell<br>organization | 3/4 | 3/4 | 4/4 | 3/3 | 3/4 | - |
| | Epidermal<br>thickness<br>[$\mu\text{m}$] | 203.3 ±<br>45.2 | 150.0 ±<br>49.7 | 128.0 ±<br>21.3 | 138.6 ±<br>35.7 | 112.2 ±<br>32.0 | 113.3 ±<br>11.9 |
|  | Collagen<br>fibers [n] | 1/4 | 3/4 | 4/4 | 3/3 | 3/4 | 3/3 |
|  | Ground<br>substance | 1/4 | 3/4 | 4/4 | 3/3 | 3/4 | 3/3 |
| | Vessels / 0.1<br>mm<sup>2</sup> | 1.5 ± 1.3 | 3.7 ± 2.3 | 6.8 ± 3.3 | 6.8 ± 2.3 | 5.8 ± 2.6 | 1.5 ± 1.2 |
| | Dermal<br>score | 2.0 ± 1.2 | 5.0 ± 1.5 | 6.2 ± 1.7 | 7.6 ± 1.5 | 6.6 ± 1.3 | 8.6 ± 0.9 |

**Table S3:** Parameters addressed to evaluate tumorigenic potential of hiPSC-derived cells. (according to published work ^44^).

|  | Parameter | Description | Outlined |
| --- | --- | --- | --- |
| **Cells** | Cell production | Cell culture research lab and expansion in CF4 | Fig. 4, 5, Fig. S7 |
|  | Quality control of cells | Cell type specific phenotype, lack of pluripotency markers | Fig. S6a |
| | Cell dose of transplanted cells | $3 \times 10^6$ KC, $3 \times 10^6$ FB, $3 \times 10^6$ ECFCs | Fig. 6, online methods |
| **Application** | Type of immunodeficient animal model | NOD.Cg-Prkdc<sup>scid</sup> Il2rg<sup>tm1Wjl</sup>/SzJ<br>B and T cell deficient; dysfunctional NK cells | https://www.jax.org/strain/005557 |
|  | Method of transplantation | Single cell suspension transplant in grafting chamber onto muscle fascia of mice, subcutaneously | Fig. 6 |
|  | Gender and numbers of animals | six female mice | online methods |
|  | Monitoring periods | 2 weeks and 12 weeks post transplantation | online methods |
| **Tumorigenesis** | Histology | Alu probing to detect human cell nuclei in the transplant | Fig. 6 |
|  |  | H&E and Ki67 showed no tumor formation | Fig. 6 |
|  | Analysis sites | Skin | Fig. 6, method section |
|  | Positive control | hiPSCs were injected into NSG mice and induced teratoma | Scharler *et al.* |

**Table S4.** List of antibodies for flow cytometry.

| Name | Host | Clonality | Clone | Working concentration | Isotype | Company | Catalog no |
| --- | --- | --- | --- | --- | --- | --- | --- |
| Tra 1-81-AF647 | mouse | monoclonal | TRA-1-81 | 60 ng/ml | IgM, k | BD Biosciences | 560793 |
| SSEA-4-PE | mouse | monoclonal | MC813-70 | 60 ng/ml | IgG3 | BD Biosciences | 560128 |
| OCT4-PE | mouse | monoclonal | 3A2A20 | 2.5 µg/ml | IgG2b, k | biolegend | 653704 |
| Nanog-PE | mouse | monoclonal | N31-355 | 78 ng/ml | IgG1, k | BD Biosciences | 560483 |
| CD56-PE | mouse | monoclonal | CMSSB | 1.25 µg/ml | IgG1, k | eBioscience | 12-0567 |
| CD90-BUV395 | mouse | monoclonal | 5E10 | 4µg/ml | IgG1, k | BD Biosciences | 563804 |
| CD105-eFluor450 | mouse | monoclonal | SN6 | 1 µg/ml | IgG1, k | eBioscience | 48-1057 |
| CD73-PE | mouse | monoclonal | AD2 | 376 ng/ml | IgG1, k | BD Biosciences | 550257 |
| CD146-PE/Vio770 | mouse | monoclonal | 541-10B2 | 1.1 µg/ml | IgG1 | Miltenyi | 130-099-956 |
| NG2-APC | mouse | monoclonal | LHM-2 | 2 µg/ml | IgG1 | R&D Systems | FAB2585A |
| CD49f-BV421 | rat | monoclonal | GoH3 | 1.25 ng/ml | IgG2k (rat) | BD Biosciences | 562582 |
| CD104-PE | rat | monoclonal | 439-9B | 1.25 ng/ml | IgG2b | BD Biosciences | 555720 |
| CD45-APC | mouse | monoclonal | HI30 | 120 ng/ml | IgG1, k | BD Biosciences | 555485 |
| CD31-eFluor450 | mouse | monoclonal | WM59 | 500 ng/ml | IgG1, k | eBiosciences | 48-0319-42 |
| Cytokeratin 14, 15, 16, 19 AF647 | mouse | monoclonal | KA4 | 2 µg/ml | IgG1 | BD Biosciences | 563648 |
| Brachyury-APC | goat | polyclonal |  | 200 ng/ml | IgG | R&D Systems | IC2085A |
| CD26-APC | mouse | monoclonal | M-A261 | 1 µg/ml | IgG1, k | BD Biosciences | 560223 |
| P63 unlabeled | mouse | monoclonal | 4E5 | 10 µg/ml | IgG1 | abcam | ab110038 |
| Cytokeratin 14 unlabeled | mouse | monoclonal | LL001 | 4 µg/ml | IgG2a | Santa Cruz | sc-53253 |
| Cytokeratin 5 unlabelled | mouse | monoclonal | 2C2 | 20 µg/ml | IgG1 | Thermo Fischer | MA5-17057 |
| anti-mouse Ig-FITC | goat | polyclonal | - | 50 µg/ml | Ig | BD Biosciences | 554001 |
| anti-mouse Ig-PE | goat | polyclonal | - | 40 µg/mL | Ig | BD Biosciences | 550589 |

## Notes

### Competing Interest Statement

Achim Schneeberger is an employee of accanis.com, a company planning to commercialize mRNA therapy.

